# Boosting Single-Cell Methods Significantly Enhances Cell-type Classification

**DOI:** 10.64898/2026.11.10.765730

**Authors:** Daniel Edelman, Helen Kang, Daniel E. Schäffer, Ekin Deniz Aksu, Bonnie Berger

## Abstract

Single-cell sequencing drives biological discovery by revealing key cellular processes, cell types, and cell states. Recent technological advances have accelerated atlas-scale data generation across laboratories. With the increasing amount of single-cell RNA-seq (scRNA-seq) data, new challenges emerge, including effective batch correction, cell-type annotation, rare cell-type and cell-state detection, as well as biological pathway discovery. Inspired by methods for boosting weak learners such as AdaBoost, here we present a novel paradigm, scBoost, that combines single-cell analysis methods using stacked ensembles to boost task performance beyond that of individual methods. We show that scBoost pretrained on Tabula Sapiens 2.0 dataset performs significantly better than existing methods in cell-type classification on the real-world Human Fetal Heart Map and Plaque Atlas. We also demonstrate that scBoost substantially improves the performance of cell-type classification and ranks at the top of the leaderboard on the OpenProblems benchmark [33], a standard-in-the-field. scBoost is modular and is adaptive to new single-cell methods, which will enable it to continue to serve as a resource to the community in the future.

**Availability:** All analysis code is made freely available at https://github.com/kanghelen/scBoost.

## Introduction

Single-cell transcriptomics technologies and analyses have accelerated biological discoveries due to their ability to detect heterogeneous cell populations, reveal new cell types and cell states, and discover causal interactions through perturbation screens [27,40,6,35,14,39,57,43,60]. To understand these high-dimensional, complex datasets, cell-type annotation becomes a crucial task in laying the foundation for many downstream analyses. Each cell type and cell state has a unique gene expression pattern, but identifying these patterns is nontrivial due to their complexity and experimental noise. Accurate and reliable cell-type annotation helps identify unique cell states and helps understand cellular heterogeneity. This will also improve downstream analyses, such as differential gene expression, marker gene discovery, cell-cell interaction, and cell state dynamics [9,19,58,59] to discover cellular pathways for potential therapeutic target development. Despite advances in annotation methods, expert annotation based on canonical marker genes remains the gold standard method [38,36]. However, this process is labor-intensive and difficult to scale, thus motivating further development of accurate, interpretable and scalable approaches for automated cell-type annotation.

Existing automated cell-type classification methods are often built around a single approach. For example, Seurat [46,21] transfer labels through anchors between mutual nearest neighbors, CellTypist [15] trains logistic regression classifiers, scANVI[56] trains a semi-supervised variational autoencoder, and Pan-human Azimuth [41] trains a neural-network classifier. Although these singleton methods have improved cell-type classification accuracy compared to previously established methods, there has not been any method that leverages the notion of a stacking ensemble from statistical machine learning to combine strengths from multiple and diverse types of single-cell embedding methods. Nearly all of these methods rely on one embedding of the cells, and embedding methods generally differ in how they weigh two opposing objectives: conserving biological variation and removing batch effects [32,33]. Published benchmarking results establish that no single method or embedding performs best across datasets and cell types [2,32,33], thus motivating the need to develop a tool that leverages multiple methods. To our knowledge, the only existing approach that combines multiple methods for scRNA-seq cell-type classification is a popular voting scheme, popV [18], which weights every method equally regardless of its accuracy. However, with popV, weaker or redundant methods can outvote stronger ones. We pursue the intuition that if we combine the embedding methods in a stacked ensemble, treating each tool as a learner, we will achieve better task performance.

Here, we present scBoost, a meta-method that combines outputs from any scRNA-seq embedding method and nominates the best combination that improves cell-type classification accuracy. scBoost is modular and generalizable to include any established or future scRNA-seq embedding methods. We utilize real-world datasets with human-annotated, gold-standard ground truth labels to train and test in cross-validation. We rigorously show that scBoost outperforms 14 published methods in OpenProblems, the recognized standard in the field for benchmarking [33]. We demonstrate that scBoost-ensemble, pretrained on Tabula Sapiens 2.0 data, performs significantly better at annotating cell types in the Human Fetal Heart Atlas [34] and the Human Plaque Atlas[50]. We demonstrate an application of agentic AI to select the optimal combination of established methods in scBoost on a given dataset. We also provide a general metric for quantifying the difference between cell embeddings. Our framework provides a universal tool for combining and evaluating scRNA-seq methods for additional scRNA-seq analysis tasks.

## Results

### Overview of scBoost

scBoost takes cell embeddings produced by any number of single-cell embedding methods as input and outputs cell-type labels. scBoost (ensemble) is a stacked-ensemble model with out-of-fold (OOF) training to prevent over-training and to enable robust cell-type classification and label transfer. scBoost (ensemble) splits the dataset into *K*-folds (here, we use *K* = 5). For each fold *k*, a base classifier is trained on one cell embedding for cells outside fold *k* and predicts cell type for cells inside fold *k*. The OOF predictions from each base learner (or meta-features) are gathered as input to *K* meta-learners. The meta-learner for fold *k* then trains on meta-features outside of fold *k*, and uses cells within fold *k* for early stopping to avoid overfitting. To prevent data leakage and to prevent overfitting, the meta-learners for one fold only see outputs from base-learners in the same fold. At inference time, the input embeddings are run through all *K*-folds’ base-learners. The base-learners’ output probabilities are concatenated and inputted to *K* meta-learners. The *K* meta-learners’ output probabilities are averaged to obtain the final prediction (Figure 1a, Methods). We contrast stacking ensemble with concatenation in scBoost-concat, which alternatively uses cell embeddings and concatenates them as input to XGBoost. In the cell-type classification task, scBoost-concat trains an XGBoost model based on concatenated cell embeddings from multiple embedding methods (Figure 1b, Methods). We also compare scBoost (ensemble) to single method embedding for cell type classification as a baseline (Figure 1c).

**Fig. 1:**
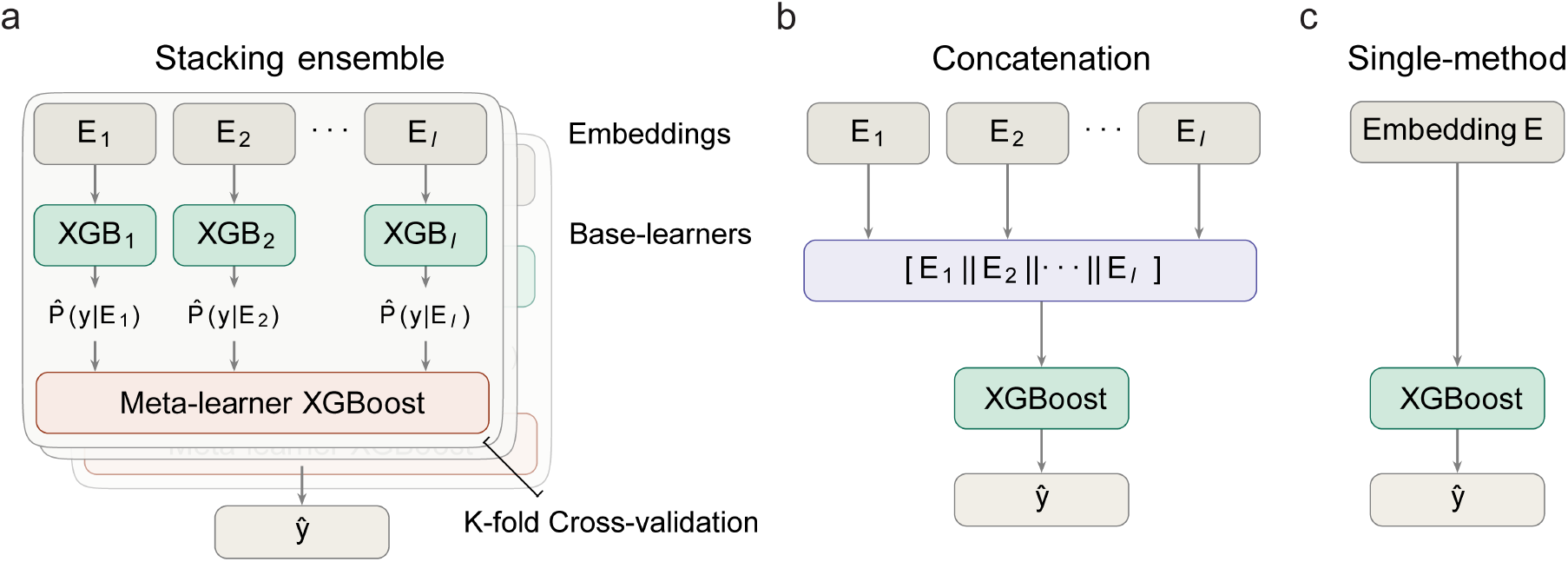
scBoost architectures for cell type classification. **a**. scBoost (ensemble) architecture. Each XGBoost base-learner is trained individually on each embeddings outputs by each of the *l* methods. **b**. scBoost-concat architecture. Concatenate *I* batch correction methods’ embeddings (scBoost-concat), then input to XGBoost cell-type classifier. **c**. Single batch correction method embeddings serve as input to XGBoost cell-type classifier.

### Pretrained scBoost model significantly outperforms existing methods in label transfer task

To test and demonstrate scBoost’s capability in label transfer, we trained scBoost (ensemble) on the Tabula Sapiens 2.0 dataset (Supplementary Methods). We applied this pretrained scBoost model to two cardiovascular datasets: Human Fetal Heart Map [34] and Human Plaque Atlas [50]. We compared scBoost to popV [18], scANVI [56], and CellTypist [15] (Supplementary Methods). In the Fetal Heart Map, scBoost out-competes popV by 37.4% (Wilcoxon signed-rank test with Holm-Bonferroni adjusted p = 4.0 × 10^*−*8^ on the donor level) in same or finer term match, and scBoost performs significantly better than all the other tested methods (Figure 2a). Additionally, scBoost performs well on all cell types. Most of the classifications on endothelial cells and endocardial cells are correct (Figure 2b,c). The next best model, popV Heart, showed less concordance with the authors’ annotation across all cell types (Figure 2b,c).

**Fig. 2:**
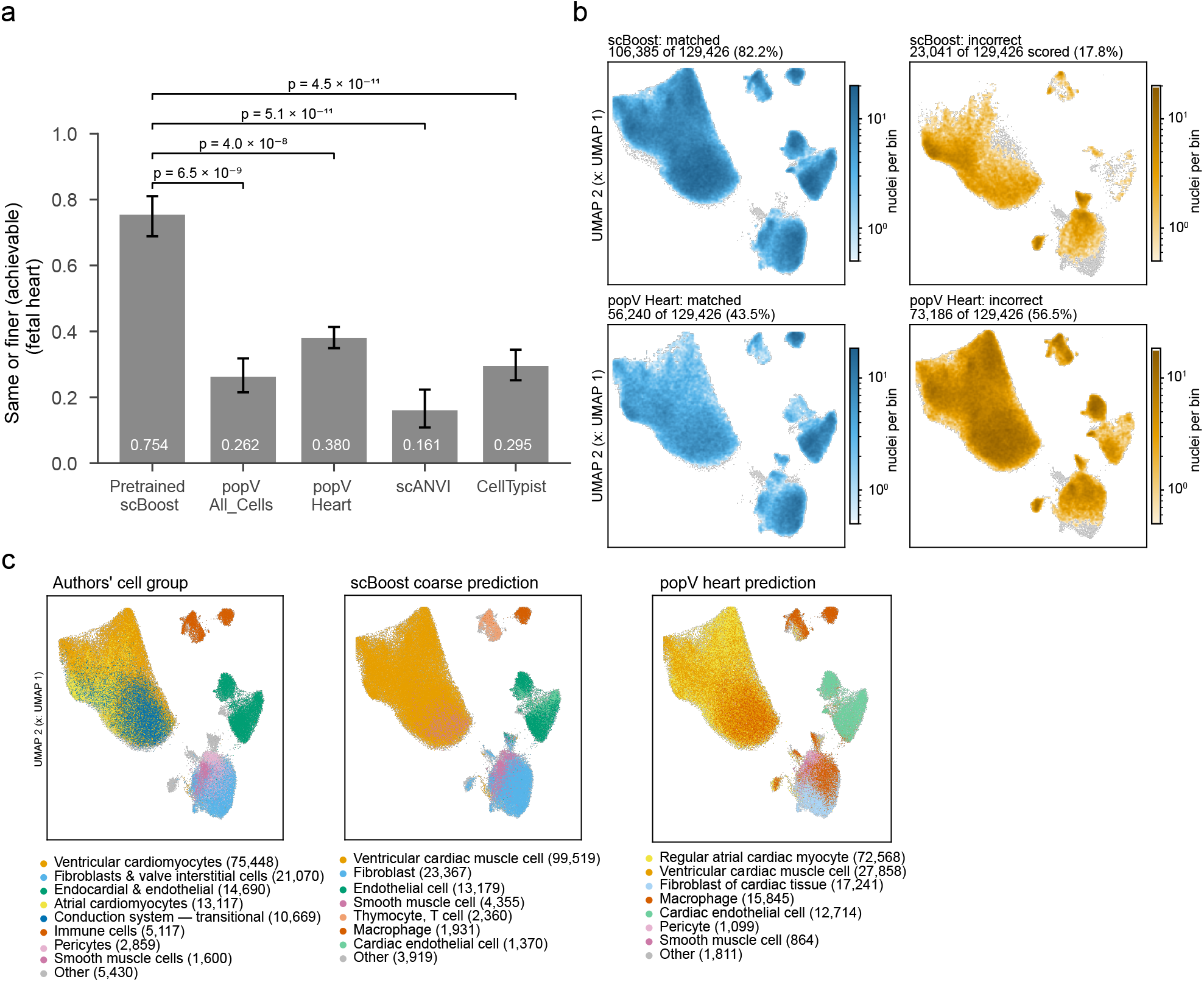
Pretrained scBoost significantly improves label transfer accuracy in the fetal heart map. **a**. The pretrained scBoost model significantly outperforms all other tested models in fetal heart map dataset. Same or finer (achievable): for Cell Ontology (CL) IDs in Tabula Sapiens 2.0, we only assess on the query cells that has a CL ID, and we count predictions with the exact term or a finer term (per Cell Ontology) as correct. **b**. UMAP displaying cells with correct or incorrect annotation. scBoost produces mostly correct annotations. popV incorrectly labels a large number of cells across all cell clusters. Matched: We count predictions with the exact term, a finer term, or a broader term per Cell Ontology as correct. **c**. UMAP comparing fetal heart map authors’ annotation, scBoost’s annotation, and popV heart pretrained model annotation. scBoost has higher concordance to the authors’ annotation than popV, especially in the lower right corner for fibroblasts and smooth muscle/mural cells.

In the Human Plaque Atlas, we also compared scBoost to 10X Genomics cellranger annotate [1], and Azimuth [41] methods. 10X Genomics cellranger (coarse) annotation produces 5,525 cells (2.8%) labeled as “cell” in the 3’ v3 chemistry dataset and 1,689 cells (2.6%) in 3’ v2 chemistry dataset (Figure S1). The Plaque Atlas provides coarse (level 1) and fine (level 2) cell type labels. We scored all models at both levels (Supplementary Methods). scBoost assigned the authors’ exact level 1 term or a finer term to 88.7% of the cells and assigns level 2 labels or a finer label per Cell Ontology [26] to 79.4% cells, which is the highest among the tested models. A per-donor statistical test shows significant accuracy improvement by scBoost compared to all other methods (Figure 3a, Tables S8, S9, Supplementary Methods). To examine annotation of transition-state cells, we investigated fibromyocytes, a modulated smooth muscle cell state discovered and characterized in Wirka et al. [53], which is also a dataset in the plaque atlas (Table S6). Fibromyocyte is a critical transitioning cell state in human plaque formation [53]. We trained scBoost on the Wirka et al. dataset and transferred its labels to the rest of the plaque atlas’s datasets. scBoost recovers 78.3% of the cells that the atlas authors call fibromyocytes, and scBoost calls a further 6,101 cells fibromyocytes that the authors labeled otherwise, specifically 3,904 labeled as fibroblasts (51% of the authors’ fibroblasts) and 2,186 labeled as smooth muscle cells (16% of the authors’ smooth muscle cells) (Figure 3b).

**Fig. 3:**
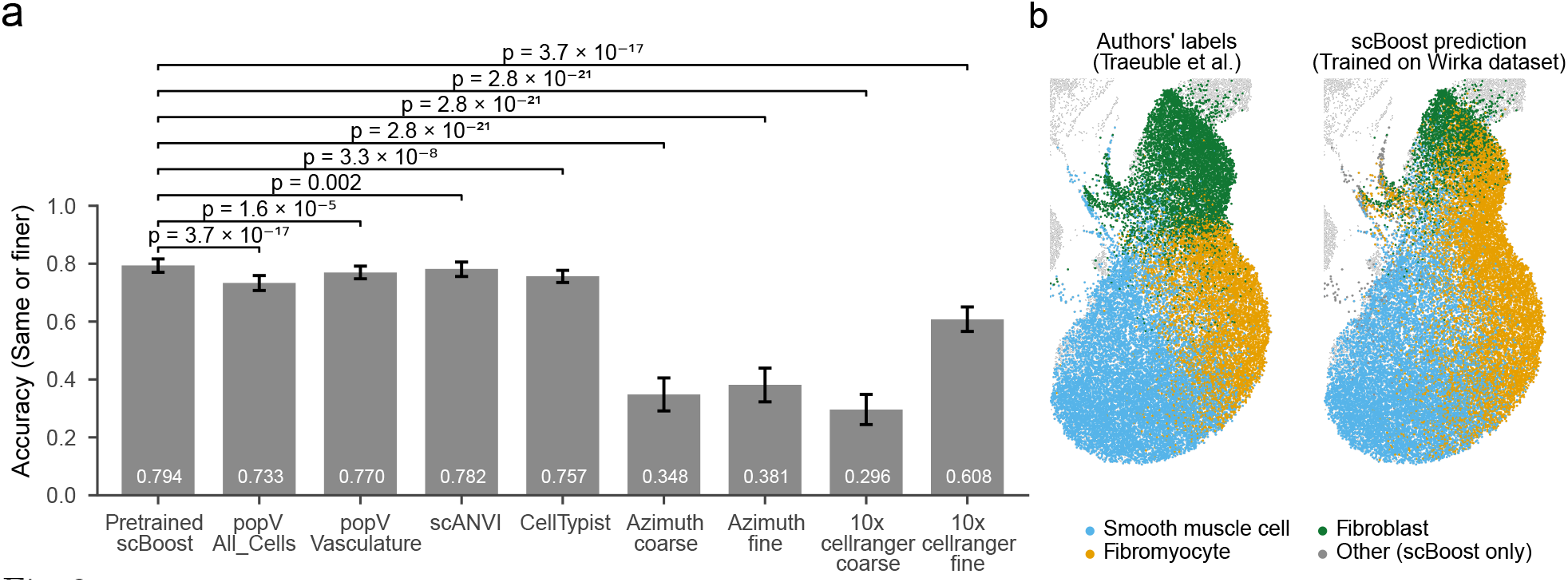
Pretrained scBoost application in the human plaque atlas. **a**. The pretrained scBoost (ensemble) model improves accuracy significantly compared to all other tested models in human plaque atlas. **b**. scBoost pretrained on the Wirka et al. dataset suggests new fibromyocyte labels. Fibromyocyte is an important activated transition state from smooth muscle cells in plaque development.

### scBoost outperforms existing methods for cell-type classification

To assess scBoost’s performance on cell type-classification, we benchmarked scBoost (ensemble) against a list of methods on the OpenProblems Platform Label Transfer task. scBoost leads in the overall score and outperforms all other published methods in OpenProblems in four out of six datasets (Figure 4a, b).

**Fig. 4:**
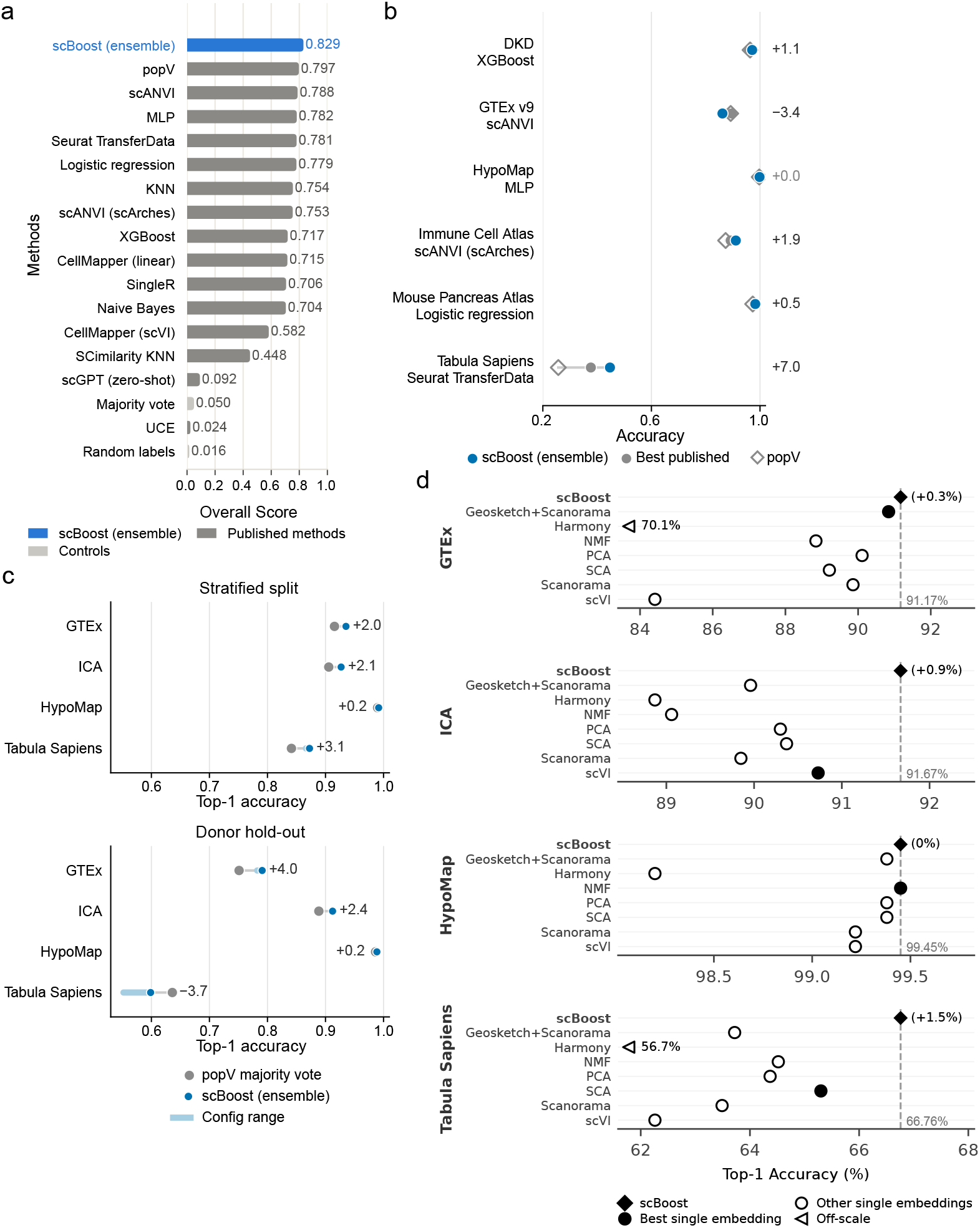
scBoost-ensemble: Stacking ensemble improves cell-type classification accuracy. **a**. scBoost-ensemble leads at the top of OpenProblems Label Transfer task score board. popV follows ranking the second. **b**. scBoost-ensemble outperforms best published methods for 4 out of 6 datasets in OpenProblems benchmarking. **c**. scBoost-ensemble and popV comparison with four input embedding methods (scVI, scANVI, Harmony, Scanorama). **d**. scBoost-ensemble performs better than single-method cell-type classification in the GTEx, ICA, and Tabula Sapiens datasets. HypoMap has saturated cell-type classification accuracy at 99.45%.

We additionally compare scBoost-ensemble against a published tool, popV[18]. popV takes a consensus among the cell type classification methods’ outputs using a popular vote as the final classification output. We add popV to OpenProblems benchmarking using popV’s default settings (Supplementary Methods). scBoost ranks at the top after this addition (Figure 4a). We note that scBoost uses a different set of embeddings than popV for this task. We decided to use popV authors’ default recommended embeddings to test on OpenProblems benchmarks, because this is what users are likely expected to use. To establish a fair comparison between popV and scBoost, we take the same set of embedding methods as input to popV and scBoost. We assess accuracy on four datasets we curated separately from OpenProblems (Supplementary Methods). scBoost performs better in top-1 accuracy compared to popV in all datasets in a stratified test setting and three out of four datasets in a donor hold-out test setting (Figure 4c). The Tabula Sapiens dataset has the most number of cell types (here, 78) and thus is challenging for both models. The substantial decrease in absolute accuracy for all holdout test results reflect the challenge in generalizing across full datasets (here, we split the dataset by donor). There is limited improvement from scBoost on the HypoMap dataset, because the models are reaching almost perfect accuracy. We note that the absolute majority of HypoMap cells are labeled as “neuron.” Overall, scBoost-ensemble achieves high cell-type classification accuracy across tested datasets.

### Single cell foundation model embeddings have a limited contribution as inputs to scBoost (ensemble)

To test whether single cell Foundation Models (scFM) embeddings improve cell-type classification with scBoost, we generate cell embeddings from scGPT, TranscriptFormer, and STATE [10,37]. To establish fair comparisons between scBoost-ensemble trained on classic and scFM embeddings, we limit scBoost to include three classical method embeddings to match the number of available scFM embeddings. scBoost-ensemble with three classic methods (1. Geosketch with Scanorama, SCA, scVI, or 2. PCA, SCA, scVI) generally outperforms those trained on scFM model embeddings (Figure 5a). To match scFM embedding dimensions with classic methods, we perform dimensionality reduction using PCA to 100 dimensions for scFM embeddings. Surprisingly, PCA dimensionality-reduced scFM model embeddings achieve higher cell-type classification accuracy than full size scFM embeddings for STATE model (Figure 5b) but not for scGPT or TranscriptFormer. This suggests dimensionality reduction applied to scFM representations for XGBoost-based cell-type classification has variable performance depending on the foundation model. To test if scFM embedding can further improve the best classic embedding combination for scBoost-ensemble, we combine scFM embedding and the best classic embedding combination (scVI + SCA) together as input. None of the scFM embeddings in addition to scVI and SCA achieve higher cell-type classification accuracy compared to scVI + SCA alone (Figure 5c). In sum, scBoost (ensemble) performs well on classic methods’ cell embeddings and scFM embeddings have limited contribution to the best classic embedding cell-type classification results.

**Fig. 5:**
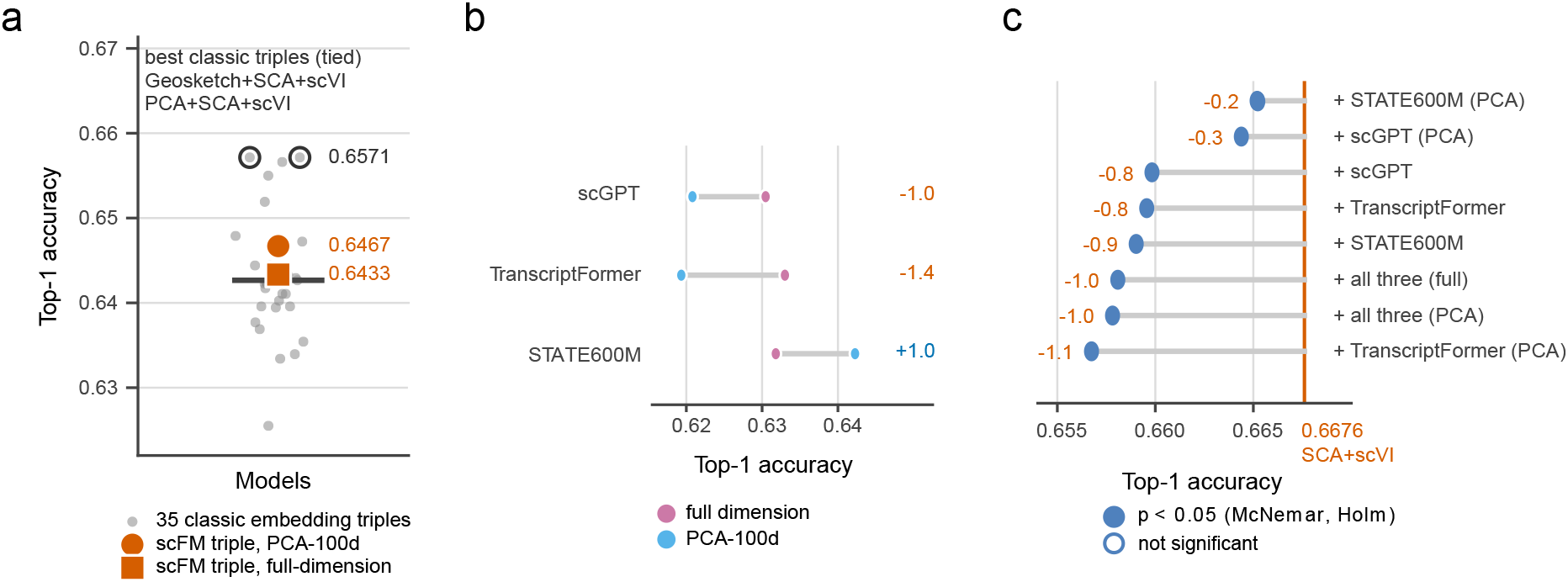
Single Cell Foundation Model Embeddings Have Limited Contribution Over Classical Cell Embeddings. **a**. Classical embedding combinations outperform scFM embedding as input to scBoost for cell type classification. To maintain fair comparisons between classic and scFM embeddings, we only show classic 3-method scBoost compared to 3-scFM embedding scBoost. **b**. Dimensionality reduction via PCA improves scBoost cell-type classification for STATE600M but not scGPT or TranscriptFormer models. **c**. scFM embeddings do not improve cell type classification accuracy by adding to the best classic embedding combinations (SCA + scVI).

### Agentic AI recommends combination of methods for scBoost

We demonstrate an application of agentic AI in recommending combinations of methods for scBoost. We sweep across all possible combination of the selected seven embedding methods, as well as scBoost’s stacking ensemble and concatenation settings. We use AI agents to analyze each dataset’s cell-type classification result, cross check the analysis, and summarize the analysis (Figure 6a). We used one agent per dataset, one agent for summarizing results, and one agent to cross-validate all results (Figure 6a). We found scBoost (ensemble) with a combination of Geosketch with Scanorama, SCA, and scVI embeddings, MLP base learners, and an XGBoost meta learner performs best in top-5 accuracy (Average top-1 accuracy: 0.8639, average top-5 accuracy: 0.9819). When focusing on top-1 accuracy, scBoost-ensemble with a combination of Geosketch with Scanorama, PCA, and Scanorama with MLP base learners and an XGBoost meta learner performs the best (Average top-1: 0.8676, average top-5: 0.9661). Furthermore, the summary agent suggests to use MLP as base-learner. Each dataset has a different top-performing combination of embedding methods (Figure 6b). The above results show that scBoost with agentic AI method selection can further improve cell-type classification accuracy and interpret parameters contributing to performance. We made the scBoost API compatible with LLM agents. In our application, agents do not choose combinations of methods for new datasets from an exhaustive search of method combinations, thus avoiding overfitting to new datasets (*e*.*g*. Fetal Heart Map and Human Plaque Atlas).

**Fig. 6:**
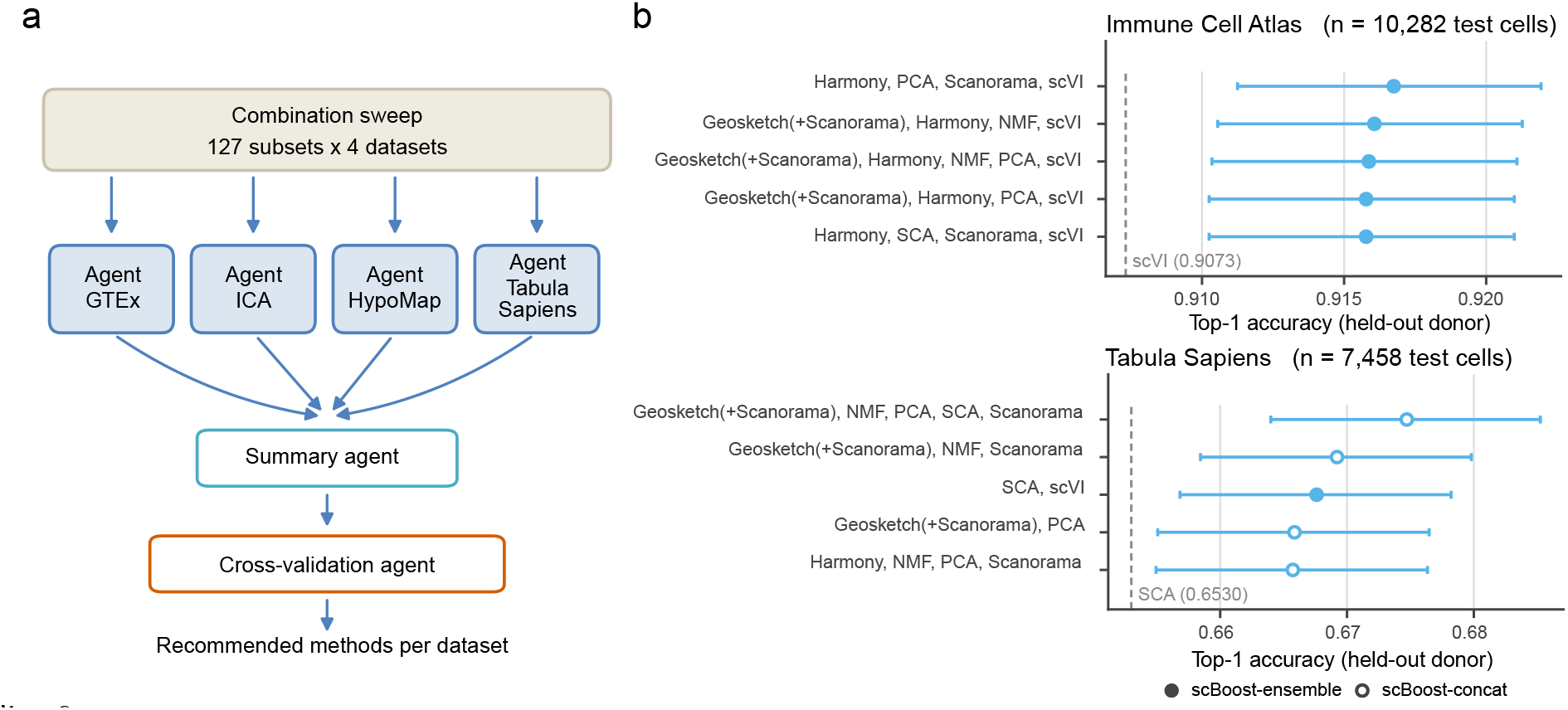
Agentic AI for selecting best combination of methods tailored to datasets. **a**. Agentic AI workflow for recommending subset of methods. **b**. Top 5 models for Immune Cell Atlas (ICA) and Tabula Sapiens datasets. Agentic AI summarizes and suggests top combination of methods for each dataset. These top 5 models have MLP base learners and an XGBoost meta learner.

### Ensembles are necessary to improve cell-type classification accuracy in scBoost

To investigate the necessity of combining multiple methods to using individual embedding methods for cell-type classification, we compare scBoost-ensemble against baseline settings with single method embeddings (1c). scBoost-ensemble generally performs better than single method cell-type classification (Figure 4d). Each dataset has a different set of top performing single embedding methods. This suggests that each method has certain strengths that work better for some datasets, and it is advantageous to combine multiple methods to draw the strengths of different methods together. In sum, scBoost improves cell type classification accuracy by combining multiple methods through a stacking ensemble approach.

### scBoost-ensemble Runtime and Memory Usage

To check the memory and time output of the scBoost, we evaluated scBoost on several datasets and with different embedding input dimensionality. We used Geosketch, Harmony, scVI, PCA, Scanorama as input to scBoost. Memory usage highly correlates with size of input data. This shows that our algorithm does not meaningfully impact the data usage (Table S11).

We evaluated time usage on 100-dimension input embeddings across all datasets that were tested for memory usage. We recorded time at various stages of scBoost processing (e.g. after data is loaded, after train/test split, after initial training, and after prediction & evaluation) (Table 1).

**Table 1:** Wall-clock runtime (seconds) of each pipeline phase across the five benchmark datasets.

|  | Runtime (s) |  |  |  |  |
|---|---|---|---|---|---|
| Phase | Tabula Sapiens | Human Immune | GTEx | HypoMap | ICA |
| Data loading | 26.7 | 3.7 | 11.1 | 22.4 | 21.6 |
| Train/test split | 3.8 | 1.3 | 2.2 | 3.0 | 2.9 |
| Fit | 7847.4 | 582.4 | 2771.9 | 171.6 | 881.9 |
| Prediction & evaluation | 14.3 | 0.8 | 9.1 | 3.4 | 12.7 |
| Total | 7904.3 | 590.2 | 2803.3 | 204.3 | 930.0 |

### Novel Metric to Quantify Embeddings’ Difference

To assess the magnitude of differences among the individual input embeddings, we constructed a metric that is the residual after point-set registration between two embeddings. We fit a linear transformation matrix between two embeddings, and the resulting loss is defined as the difference between two embeddings. The problem can be viewed as finding a transition matrix *T* such that *TA* = *B*, where *A, B* are embedding matrices for cells where the same rows in each have the same cells. The distance would be the resulting norm of the residuals ∥*TA* −*B*∥ _2_. To prevent issues of scale confounding the loss, the embeddings are first normalized and centered. We created two methods for this calculation: 1) an iterative solver attempting to calculate loss by taking embedding pairs for batches of cells, and 2) a linear least-square solver.

We computed the distance between embeddings for across four datasets (Tabula Sapiens, GTEx, Immune Cell Atlas and HypoMap) and five methods (GeoSketch + Scanorama, Harmony, PCA, Scanorama and scVI) (Figure 7). PCA and Harmony cluster together and Geosketch + Scanorama clusters together with Scanorama. We can utilize this information to select for methods that have high distances from each other as a first try in scBoost. This narrows the search space for scBoost and could thereby reduce overfitting that would arise from an exhaustive search over similar or related embedding methods.

**Fig. 7:**
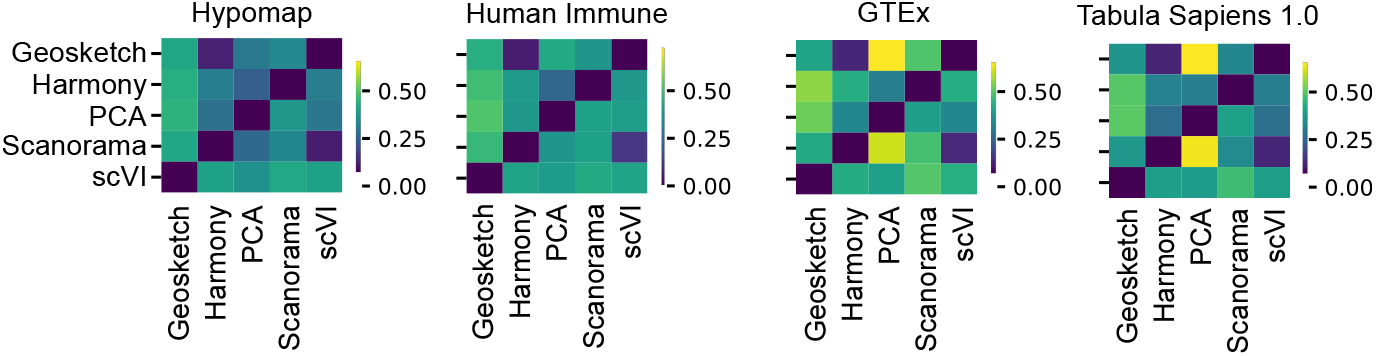
Heatmap of comparison of methods, linear least square solver.

## Discussion

scRNA-seq technologies enable measurement of high-throughput gene expression and detection of multiple cell types and states simultaneously. Identifying cell types accurately can set a solid foundation for downstream tasks. Numerous scRNA-seq analysis methods are designed to perform various tasks and have unique advantages and limitations. Benchmarking studies such as the OpenProblems [33] platform compare the performances of these methods. Despite one study that combined scRNA-seq analysis methods via popular vote [18], no existing tool adapts stacking ensemble across different cell embeddings methods for scRNA-seq to further boost cell-type classification performance.

Our study introduces a new meta-method, where we combine outputs from various established scRNA-seq embedding methods and, in turn, improve cell type classification performance compared to any of the methods alone. While many previous methods utilize boosting on a single method, or take the consensus from a set of methods, we are the first to propose an architecture that combines multiple single cell methods for downstream tasks (scBoost-concat) and utilizes stacking ensemble (scBoost). We select methods from the scverse, and we select methods base on past benchmarking studies [22,24,32,33], but any scRNA-seq cell embedding methods will be suitable for scBoost. sc-Boost (ensemble) trains separate base classifiers on each method’s embedding and combines their probabilistic outputs through a meta-learner. Stacking enables scBoost to automatically discover which embedding methods are most informative for which cell types to improve cell-type classification accuracy. Our out-of-fold training for both base-learners and meta-learners ensures no data leakage and prevents overfitting.

We applied pretrained scBoost to the Fetal Heart Map and Human Plaque Atlas datasets, and showed that scBoost significantly outperforms other tested methods. To demonstrate scBoost’s capability to identify a critical transition state in disease development, we trained scBoost on human coronary artery scRNA-seq data (Wirka et al. [53]) with the dataset authors’ gold standard labels and use this scBoost model to suggest new labeling for human plaque atlas on fibromyocyte, which is an important modulated smooth muscle cell state in human plaque development. We also benchmarked scBoost on the established OpenProblems platform [33] for the label transfer task and showed that scBoost ranks at the top of the scoreboard.

To nominate the best combination of embedding methods for each dataset, we created an embedding method difference metric. We note that this metric is not commutative and therefore is not called a distance metric between cell embeddings. We also made the scBoost API compatible with LLM agents and demonstrated that we can leverage agentic AI to suggest best combination for scBoost in a dataset-specific way.

Learned combinations of existing methods have also improved other genomics tasks. For example, pgBoost trains a gradient-boosted model on the scores of single-cell multiome linking methods and genomic distance, and links regulatory variants to target genes better than any of these methods alone [16]. pgBoost and scBoost-concat share a similar architecture, where one boosted model operates on concatenated method outputs. In contrast, scBoost (ensemble) trains one classifier per method and stacks their out-of-fold predictions. Supervised combinations of existing methods and features have also improved enhancer-gene linking from single-cell multiome data [44]. We envision that scBoost will improve performance for additional single cell genomics and analysis tasks such as Perturb-seq prediction, multimodal label transfer, cross-species transfer, and gene prioritization.

In sum, scBoost is a novel scRNA-seq cell type classification method, and it can be considered as a new framework that leverages ideas from statistical boosting to improve scRNA-seq analysis performance. Our framework is broadly generalizable to any scRNA-seq analysis tasks beyond cell-type classification. We anticipate scBoost to be a valuable resource and to serve as a foundation for further improving scRNA-seq analysis methods.

## Methods

### scBoost Architecture

#### scBoost (ensemble)

##### Prior work

Statistical boosting algorithms have been theoretically and empirically shown to leverage weak learners to improve task performance [42]. Ensembles of a single type of learner have been applied to single-cell analysis for cell-type classification (*e*.*g*., XGBoost models [8,3,51], scClassify [30]), gene regulatory network inference (*e*.*g*., GENIE3 [25], GRNBoost [5] and XGRN [13]) and marker gene discovery (*e*.*g*., EDGE [47] and NS-Forest [31]). On the other hand, stacking [55,7] instead trains a meta-learner on held-out predictions of diverse base learners; thus, the model learns how much to trust each of the base learners. It has been shown that with cross-validation, stacking performs asymptotically as well as the best of its candidate combinations [29].

To combine embeddings for cell-type classification, we consider a two-level stacking ensemble model [11] (Supplementary Algorithm S2, Fig.1a). Rather than operating in the raw gene expression space, stacking trains a separate base classifier for each method embedding and a meta-learner to combine all base-learner’s probabilistic outputs.

##### Stage 1: Base classifiers with out-of-fold predictions

For each method *i*, we train a multi-layer perceptron (MLP) base classifier *f*_*i*_ on embedding *E*_*i*_ using *K*-fold cross-validation. Here, we select *K* = 5. For each fold *k*, the classifier 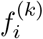 is trained on cells outside fold *k* and produces class probability predictions 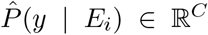 for the held-out cells, where *C* is the number of cell-types. This out-of-fold (OOF) scheme prevents information leakage from base classifiers to the meta-learner. After all folds, every training cell has an OOF probability vector from each method, assembled into a meta-feature matrix for *I* methods, *n*_*train*_ cells, and *C* cell types:

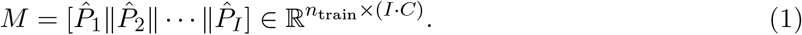

##### Stage 2: Meta-learner

We next fit *K* = 5 meta-learners using the same fold assignment in Stage 1. Meta-learner *g*_*k*_ is trained on the rows of *M* belonging to cells outside fold *k*, while the rows inside fold *k* are held out as its validation set for early stopping. To learn which base classifiers to trust for which cell-types, each *g*_*k*_ operates in the *I* methods × *C* cell-type dimensional probability space. To prevent leakage, we make fold *k* invisible to meta-learner *g*_*k*_. With donor-grouped folds, no donor is shared between the training and validation sets of any *g*_*k*_.

The choice of meta-learner family is independent of the base learner training. We selected gradient-boosted trees (XGBoost) as the default meta-learner, and an elastic-net multinomial logistic regression as an alternative. In both cases, we use the held-out fold for early stopping.

##### Stage 3: Test-time prediction

Prediction proceeds through *K* sets of base learners 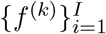 and their corresponding meta-learner *g*_*k*_ trained on the same folds. We pass test cells through each set of base classifiers and concatenate their probabilities into

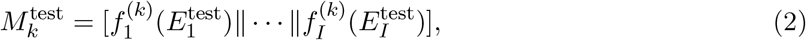

Then, the matching meta-learner *g*_*k*_ scores on 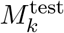. The final prediction averages the outputs from *K* meta-learners and selects for the cell-type *c* with the highest probability:

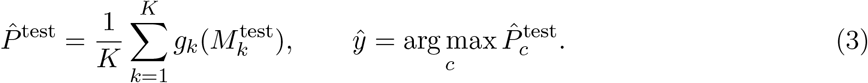

#### scBoost-concat

To combine embeddings from *I* methods, we first concatenate them into a single feature matrix and optionally reduce it via principal component analysis (PCA) or surprisal component analysis (SCA) [12] to a target dimension *d* before classification. Classification proceeds with using an XGBoost classifier (Figure 1b, Algorithm S1).

## Supplementary Material

### scBoost algorithm

#### Algorithm S1

scBoost-concat: Combining Batch Correction Embeddings

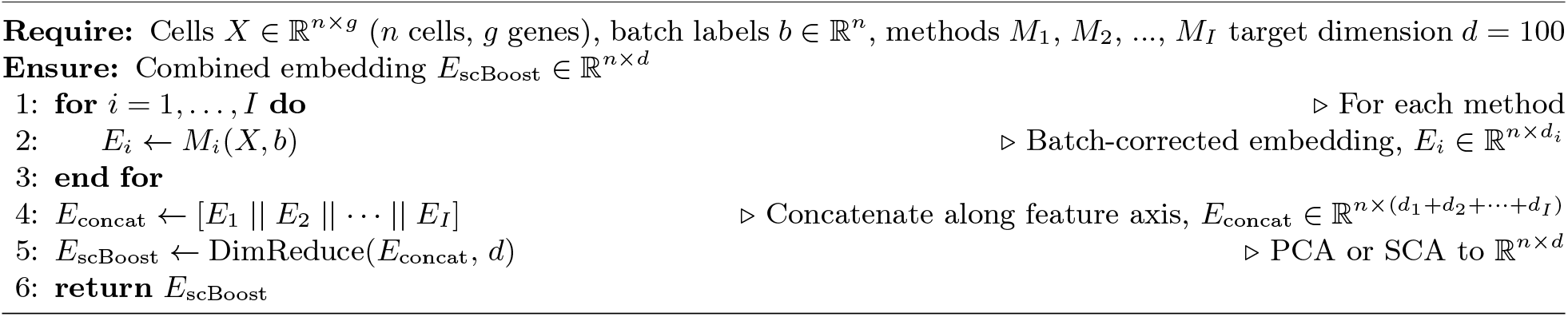

#### Algorithm S2

scBoost-ensemble with out-of-fold predictions

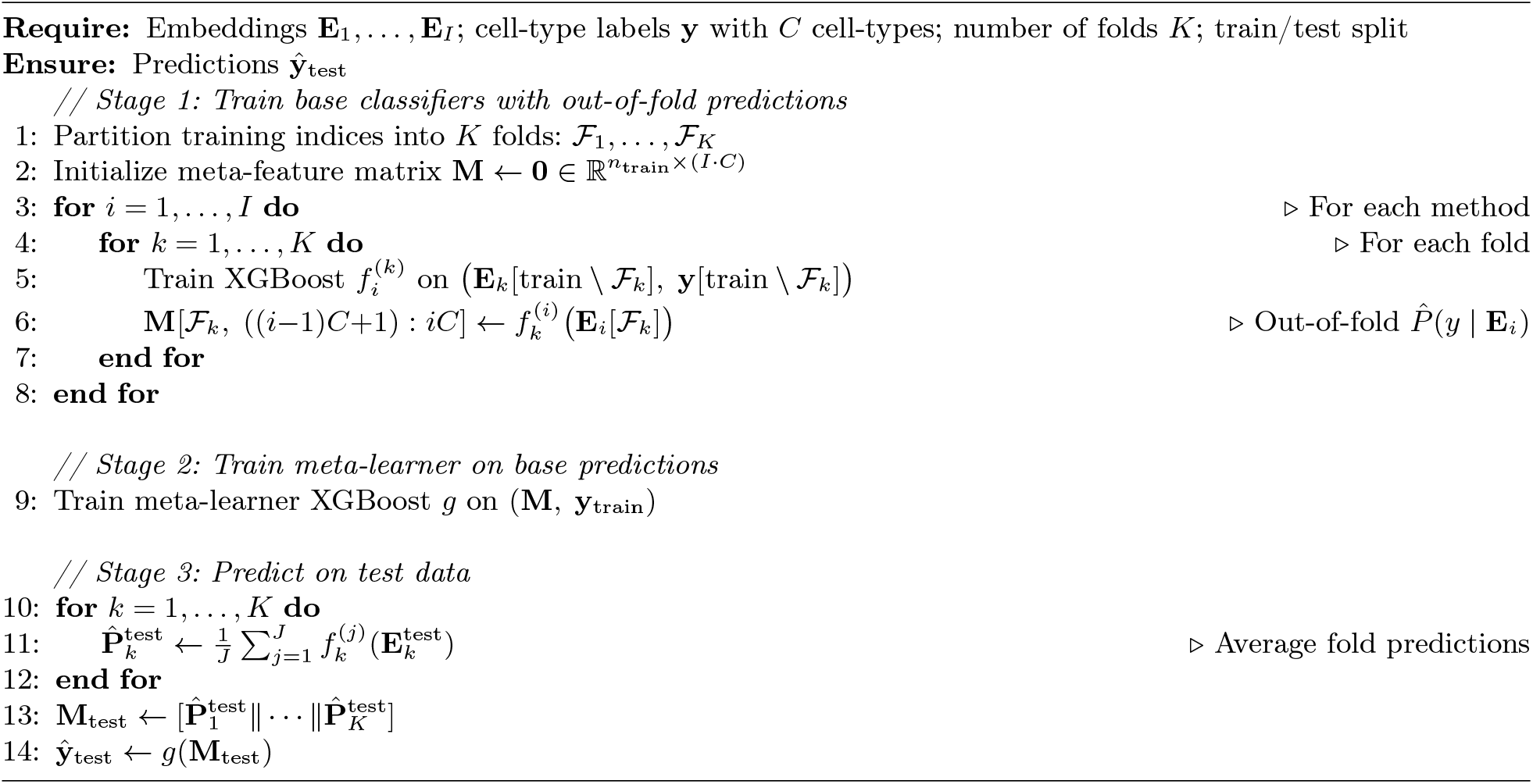

### Single-Cell RNA-seq Datasets for scBoost benchmarking

We utilized publicly available datasets (Supplementary Table S2): Tabula Sapiens [49] (the first draft of the human cell atlas), GTEx data, Human Immune Cell Atlas (ICA), and Hypomap. All datasets have GeoSketch [23] sampling to 100,000 cells. For stratified split experiments, we split the cells into 90% training and 10% test datasets stratified by cell type to ensure the model will see all cell types in the same ratio as the input data. For experiments splitting datasets with **holdout** data, we select one or more donors that has a sum of cells that is the closest to 10% data as the holdout set. This prevents data leakage and mimics having a brand new dataset for cell-type classification. For methods that do not require training, we use the same 10% test dataset for evaluating cell-type annotation accuracy to ensure consistency.

Tabula Sapiens 1.0 dataset has many fine labeled cells, with a total of 180 cell types, that have low cell count per cell type. Therefore, we manually annotated the cell types and used this new label set of 78 cell types in training scBoost and benchmarking (Supplementary Table S1).

**Table S1:** Tabula Sapiens cell types updated for the benchmark: the authors’ 160 Cell Ontology types grouped into 79 updated labels. Cells are counted in the CELLxGENE release (483,152 cells) and in the 100,000-cell benchmark subsample; Sperm does not occur in the subsample.

| **Updated Label** | **Author Cell Types (Cell Ontology)** | **Cells,**<br>**Release** | **Cells,**<br>**Subsample** |
|---|---|---|---|
| Acinar cell | acinar cell of salivary gland, pancreatic acinar cell | 15,681 | 1,335 |
| Adipocyte | fat cell | 44 | 33 |
| Adventitial cell | adventitial cell | 815 | 323 |
| B cell | B cell | 19,807 | 1,628 |
| Basal cell | basal cell | 14,065 | 2,702 |
| Basophil | basophil | 715 | 138 |
| CD4<sup>+</sup> T cell | CD4-positive, alpha-beta T cell, CD4-positive helper T cell, effector CD4-positive, alpha-beta T cell, CD4-positive, alpha-beta memory T cell, naive thymus-derived CD4-positive, alpha-beta T cell, T follicular helper cell | 38,297 | 4,464 |
| CD8<sup>+</sup> T cell | CD8-positive, alpha-beta T cell, CD8-positive, alpha-beta memory T cell, effector CD8-positive, alpha-beta T cell, CD8-positive, alpha-beta cytokine secreting effector T cell, naive thymus-derived CD8-positive, alpha-beta T cell, CD8-positive, alpha-beta cytotoxic T cell | 26,975 | 4,605 |
| Cardiac muscle cell | cardiac muscle cell | 7,205 | 439 |
| Ciliated epithelial cell | lung ciliated cell, ciliated cell, ciliated epithelial cell | 950 | 626 |
| Club cell | club cell | 2,116 | 656 |
| Conjunctival epithelial cell | conjunctival epithelial cell | 4,587 | 1,477 |
| Connective tissue cell | connective tissue cell, tendon cell | 483 | 154 |
| Corneal epithelial cell | corneal epithelial cell | 1,488 | 323 |
| Dendritic cell | dendritic cell, plasmacytoid dendritic cell, Langerhans cell, liver dendritic cell, mature conventional dendritic cell | 716 | 230 |
| Duct epithelial cell | duct epithelial cell | 1,693 | 534 |
| Endothelial cell | endothelial cell, cardiac endothelial cell, endothelial cell of lymphatic vessel, lung microvascular endothelial cell, endothelial cell of hepatic sinusoid, gut endothelial cell | 17,906 | 5,923 |
| Enterocyte | enterocyte of epithelium of large intestine, enterocyte, enterocyte of epithelium of small intestine | 6,270 | 1,134 |
| Enteroendocrine cell | intestinal enteroendocrine cell | 113 | 59 |
| Epithelial cell | epithelial cell, myoepithelial cell, epithelial cell of uterus, medullary thymic epithelial cell, surface ectodermal cell, epithelial cell of lacrimal sac, salivary gland cell, duodenum glandular cell, intrahepatic cholangiocyte, pigmented ciliary epithelial cell | 9,972 | 1,476 |
| Erythroid cell | erythrocyte, erythroid progenitor cell, erythroid lineage cell | 11,877 | 744 |
| Eye photoreceptor cell | eye photoreceptor cell | 891 | 294 |
| Fibroblast | fibroblast, fibroblast of breast, fibroblast of cardiac tissue | 34,639 | 9,302 |
| Glial cell | microglial cell, radial glial cell | 404 | 226 |
| Goblet cell | large intestine goblet cell, goblet cell, small intestine goblet cell | 996 | 382 |
| Granulocyte | granulocyte | 855 | 440 |
| Hematopoietic stem cell | hematopoietic stem cell | 794 | 424 |
| Hepatocyte | hepatocyte | 2,647 | 761 |
| Innate lymphoid cell | innate lymphoid cell | 5,131 | 681 |
| Intestinal stem cell | intestinal crypt stem cell, intestinal crypt stem cell of large intestine, intestinal crypt stem cell of small intestine | 430 | 153 |
| Intestinal tuft cell | intestinal tuft cell | 193 | 82 |
| Ionocyte | ionocyte, pulmonary ionocyte | 119 | 79 |
| Keratinocyte | keratinocyte | 3,667 | 251 |
| Keratocyte | keratocyte | 595 | 239 |
| Kidney epithelial cell | kidney epithelial cell | 8,331 | 438 |
| Leukocyte | leukocyte | 2,286 | 849 |
| Luminal epithelial cell | luminal epithelial cell of mammary gland | 5,378 | 1,361 |
| Macrophage | macrophage | 35,204 | 9,121 |
| Mast cell | mast cell | 2,238 | 1,097 |
| Melanocyte | melanocyte | 208 | 90 |
| Memory B cell | memory B cell | 15,676 | 1,234 |
| Mesenchymal stem cell | mesenchymal stem cell | 15,459 | 2,574 |
| Mesothelial cell | mesothelial cell | 41 | 21 |
| Monocyte | classical monocyte, monocyte, non-classical monocyte, intermediate monocyte | 27,634 | 7,556 |
| Muscle cell | muscle cell | 437 | 198 |
| Myeloid cell | myeloid cell | 2,227 | 1,037 |
| Myeloid dendritic cell | CD1c-positive myeloid dendritic cell, CD141-positive myeloid dendritic cell, myeloid dendritic cell | 493 | 292 |
| Myeloid progenitor | common myeloid progenitor | 290 | 121 |
| Myofibroblast | myofibroblast cell | 3,070 | 679 |
| Müller cell | Mueller cell | 360 | 211 |
| NK/T cell | mature NK T cell, type I NK T cell, immature natural killer cell | 15,697 | 3,233 |
| Naive B cell | naive B cell | 6,761 | 285 |
| Neutrophil | neutrophil | 17,694 | 2,227 |
| Pancreatic ductal cell | pancreatic ductal cell | 2,431 | 827 |
| Pancreatic endocrine cell | type B pancreatic cell, pancreatic PP cell, pancreatic A cell, pancreatic D cell | 250 | 109 |
| Pancreatic stellate cell | pancreatic stellate cell | 393 | 151 |
| Paneth cell | paneth cell of colon, paneth cell of epithelium of small intestine | 1,535 | 407 |
| Pericyte | pericyte | 4,539 | 1,565 |
| Plasma cell | plasma cell, plasmablast | 11,559 | 4,124 |
| Platelet | platelet | 268 | 174 |
| Pneumocyte | type II pneumocyte, type I pneumocyte | 9,357 | 965 |
| Prostate epithelium | basal cell of prostate epithelium, luminal cell of prostate epithelium | 4,390 | 1,404 |
| Respiratory goblet cell | respiratory goblet cell, tracheal goblet cell | 839 | 341 |
| Retinal neuron | retinal bipolar neuron, retina horizontal cell, retinal ganglion cell | 45 | 37 |
| Retinal pigment epithelial cell | retinal pigment epithelial cell | 49 | 33 |
| Schwann cell | Schwann cell | 10 | 7 |
| Secretory epithelial cell | secretory cell, mucus secreting cell, serous cell of epithelium of trachea, serous cell of epithelium of bronchus | 373 | 183 |
| Skeletal muscle cell | fast muscle cell, slow muscle cell, tongue muscle cell, cell of skeletal muscle | 585 | 394 |
| Skeletal muscle satellite cell | skeletal muscle satellite stem cell | 4,831 | 1,398 |
| Smooth muscle cell | smooth muscle cell, vascular associated smooth muscle cell, myometrial cell, bronchial smooth muscle cell | 7,031 | 2,732 |
| Sperm | sperm | 11 | 0 |
| Stem cell | stem cell | 144 | 92 |
| Stromal cell | stromal cell | 3,411 | 898 |
| T cell | T cell | 14,682 | 1,913 |
| Thymocyte | DN3 thymocyte, DN1 thymic pro-T cell, thymocyte, DN4 thymocyte, double-positive, alpha-beta thymocyte | 3,364 | 290 |
| Transit amplifying cell | transit amplifying cell of small intestine, transit amplifying cell of colon | 435 | 130 |
| Treg cell | regulatory T cell, naive regulatory T cell | 2,064 | 526 |
| Urothelial cell | bladder urothelial cell | 4,151 | 1,486 |
| Vascular endothelial cell | capillary endothelial cell, endothelial cell of vascular tree, vein endothelial cell, endothelial cell of artery, retinal blood vessel endothelial cell, blood vessel endothelial cell | 13,785 | 4,873 |

For benchmarking with the OpenProblems pipeline, we downloaded six datasets that were used to benchmark all other published methods: diabetic kidney disease (DKD), GTEx v9, HypoMap, Immune Cell Atlas, Mouse Pancreas Atlas, and Tabula Sapiens.

For scBoost application on cardiac datasets, we downloaded the Human Fetal Heart Map [34] and the Human Plaque Atlas [50].

**Table S2:**
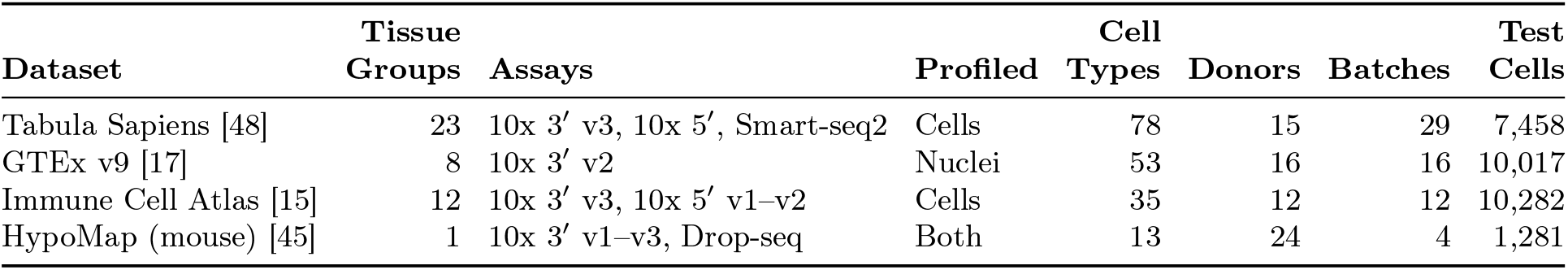
Benchmark datasets, each are subsampled to 100,000-cell via Geosketch. Tabula Sapiens, GTEx, and Immune Cell Atlas are human samples. Tissue groups: the number of unique CELLx-GENE’s general tissue terms. Test cells: the number of cells from one held-out donor.

|  | **Tissue** |  |  | **Cell** |  |  | **Test** |
|---|---|---|---|---|---|---|---|
| **Dataset** | **Groups** | **Assays** | **Profiled** | **Types** | **Donors** | **Batches** | **Cells** |
| Tabula Sapiens [48] | 23 | 10x 3′ v3, 10x 5′, Smart-seq2 | Cells | 78 | 15 | 29 | 7,458 |
| GTEx v9 [17] | 8 | 10x 3′ v2 | Nuclei | 53 | 16 | 16 | 10,017 |
| Immune Cell Atlas [15] | 12 | 10x 3′ v3, 10x 5′ v1–v2 | Cells | 35 | 12 | 12 | 10,282 |
| HypoMap (mouse) [45] | 1 | 10x 3′ v1–v3, Drop-seq | Both | 13 | 24 | 4 | 1,281 |

### Generating Cell Embeddings

We selected commonly-used scRNA-seq embedding methods: Scanorama [22,24], Scanorama with Geosketch [23], Harmony [28], scVI [20,52], SCA [12], NMF, and PCA. We set the dimension for each output embedding to be 100. For single-cell foundation models, we tested scGPT [10], STATE [4] and Transcriptformer [37]. We used the default settings for each model and set the output embedding dimension to the default embedding dimension: scGPT to 512, STATE to 2058, and Transcriptformer to 2048. To perform a fair comparison between cell embeddings generated from individual methods and scBoost embeddings, we used Principal Component Analysis (PCA) [54] to reduce the final embedding dimension to 100 prior to evaluation.

### Training Dataset, Architecture, and Cell Ontology Classes for scBoost Pretrained Model

We used the Tabula Sapiens 2.0 (CELLxGENE Tabula Sapiens - All Cells [49]) dataset for training scBoost-ensemble. To allow for label transfer, we selected embedding methods that allows for data projection without refitting: PCA (100 dimensions from 3,000 highly variable genes (HVGs)), NMF (100 dimensions from 3,000 HVGs), scVI (30 dimensions from 2,000 HVGs). The Tabula Spiens 2.0 datasets are labeled with Cell Ontology (CL) terms. We trained scBoost-ensemble based on these CL labels. For CL terms with less than 20 cells, we assign these cells into their nearest super-set within two steps that also appeared in the training dataset, otherwise the cells are set aside (Supplementary Table S3). For CL terms that only exists in one donor, we randomly spread the cells in those CL terms across folds such that every fold’s base classifiers see the term (Supplementary Table S4).

### Evaluation Metrics

We primarily used top-1 accuracy to assess cell-type classification accuracy. We also report top-5 accuracy when applicable.

**Table S3:** Tabula Sapiens 2.0 label assignment. There are 13 labels spanning only 94 cells. A label is merged into the nearest trained ancestor at most two steps up in the Cell Ontology (release v2026-06-08). Otherwise, the cells are excluded from training (assigned to none in the table).

| **Label (Tabula Sapiens v2)** | **CL ID** | **# Cells** | **# Donors** | **Assigned class** |
|---|---|---|---|---|
| **Merged into the nearest trained ancestor** |  |  |  |  |
| radial glial cell | CL:0000681 | 13 | 3 | glial cell |
| pancreatic D cell | CL:0000173 | 9 | 1 | enteroendocrine cell |
| T follicular helper cell | CL:0002038 | 6 | 1 | CD4-positive, alpha-beta T cell |
| pancreatic PP cell | CL:0002275 | 2 | 1 | enteroendocrine cell |
| erythroid lineage cell | CL:0000764 | 2 | 1 | myeloid cell |
| retinal ganglion cell | CL:0000740 | 1 | 1 | neuron |
| **Set aside (no trained ancestor within two steps)** |  |  |  |  |
| oocyte | CL:0000023 | 18 | 1 | none |
| serous cell of epithelium of bronchus | CL:1000331 | 15 | 2 | none |
| Leydig cell | CL:0000178 | 10 | 1 | none |
| unknown | — | 9 | 1 | none |
| retina horizontal cell | CL:0000745 | 5 | 2 | none |
| epithelial fate stem cell | CL:0000036 | 2 | 1 | none |
| mesenchymal cell | CL:0008019 | 2 | 1 | none |

**Table S4:** Trained classes with a single donor. Nine of the 167 trained Tabula Sapiens 2.0 classes have cells from one donor (TSP1, TSP2, TSP25, TSP27).

| **Class** | **CL ID** | **Compartment** | **Tissue** | **Donor** | **Cells** | **Trained** |
|---|---|---|---|---|---|---|
| kidney epithelial cell | CL:0002518 | Epithelium | kidney | TSP2 | 9,278 | 3,000 |
| spermatid | CL:0000018 | Germline | testis | TSP25 | 4,009 | 3,000 |
| spermatocyte | CL:0000017 | Germline | testis | TSP25 | 3,070 | 3,000 |
| enterocyte of epithelium proper of jejunum | CL:1000341 | Epithelium | small intestine | TSP2 | 405 | 405 |
| male germ cell | CL:0000015 | Germline | testis | TSP25 | 116 | 116 |
| interstitial cell of Cajal | CL:0002088 | Epithelium | stomach | TSP27 | 71 | 71 |
| peritubular myoid cell | CL:0002481 | Epithelium | testis | TSP25 | 67 | 67 |
| spermatogonium | CL:0000020 | Germline | testis | TSP25 | 64 | 64 |
| pancreatic A cell | CL:0000171 | Epithelium | pancreas | TSP1 | 48 | 48 |

For OpenProblems benchmarking, we used the same metrics considered in the OpenProblems pipeline: accuracy, F1 macro, F1 micro, and F1 weighted scores.

For pretrained scBoost (ensemble) predictions on cardiac datasets, we constructed four criteria for measuring cell type classification accuracy:

#### Exact

the prediction is the true term. It is computed on the cells whose true term is one of the 167 Tabula Sapiens 2.0 classes.

#### Same or finer

the prediction is the true term or a finer term below it in Cell Ontology. A correct subtype call counts but a broader term prediction does not count as correct.

#### Same or finer (achievable)

Same or finer, counted only on the cells whose true term has a same or finer match among the Tabula Sapiens 2.0 classes; the other cells are left out.

#### Same, finer or coarser

the prediction is the true term, a finer term or a broader one per Cell Ontology. In other words, any call on the cell’s lineage counts as correct.

### Class prior of the reference-trained model

The reference-trained scBoost (ensemble) model is trained on *N* = 637,267 Tabula Sapiens 2.0 cells with embeddings *x*_*i*_ and labels *y*_*i*_ ∈{1,…,*K}, K* = 167 classes. Class *c* has *n*_*c*_ training cells and the share *π*_*c*_ = *n*_*c*_*/N* of the training set, its training prior. Each of the model’s learners (15 base classifiers (3 embeddings: PCA, NMF, scVI, and 5 folds) and 5 meta-learners) minimizes the mean cross-entropy

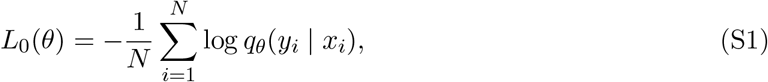

where *q*_*θ*_(*c x*) is the posterior the learner outputs. Because every cell counts once, a class with *n*_*c*_ cells contributes *n*_*c*_ terms. Therefore, the minimizer is the Bayes posterior of the training set,

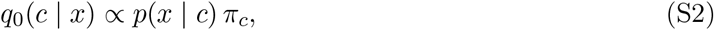

where *p*(*x* | *c*) is the fit of embedding *x* to class *c*. The call is the fit multiplied by the label’s training set size. In other words, a class that fits a cell slightly worse will have high prediction probability if *π*_*c*_ large enough. In Tabula Sapiens 2.0, the parent terms are larger than their subtypes, and scBoost (ensemble) in term label some tissues subtype only by its parent cell type.

We construct a class prior *β* to remove the size advantage from the loss. A training cell of class *c* carries the weight

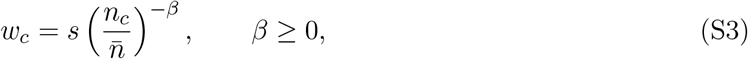

where 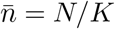 and *s* sets the mean weight per cell to one, and each learner minimizes the weighted cross-entropy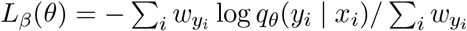, for training and for early stopping. At *β* = 0 every weight is one and *L*_*β*_ = *L*_0_; at *β* = 1 the total weight *n*_*c*_*w*_*c*_ is the same for every class, so each class pulls on the model equally whatever its size. With enough data the minimizer of *L*_*β*_ is

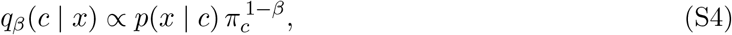

and the prior is now 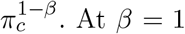, equation S4 has a flat prior. Therefore, when two classes (*a* and *b*) are imbalanced but class *a* has a higher probability, regardless of class size, the model will still call class *a* over class *b*:

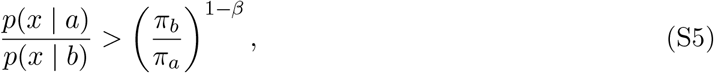

The right hand side is a likelihood-ratio threshold. The threshold is the full prior ratio at *β* = 0 and 1 (or flat) at *β* = 1. At *β <* 1, this threshold protects large classes, because if a cell from a large class has a higher prediction probability in a smaller class by chance, this cell can still receive the correct prediction.

We fit models at *β* = 0, 0.5, and 1 to allow for trade-off between coarse and fine calls. Like 10x cellranger annotate and Azimuth methods, scBoost can have the option to give coarse annotations and fine annotations.

### Label transfer with scBoost-ensemble for the Human Plaque Atlas and Fetal Heart Atlas

We trained scBoost (ensemble) on the Tabula Sapiens 2.0 dataset with the authors’ Cell Ontology (CL) labels. We used two-hidden-layer MLP base learners and tested two versions of meta-learners (Logistic Regression, XGBoost). We default to MLP base learners, XGBoost meta-learner, and *β* = 0 for the main section results, and we include all results in the Supplementary Tables.

The Human Plaque Atlas contains cells collected via different 10x Genomics assay versions. Some cells are sorted for CD45 marker and some cells are unsorted. The Human Plaque Atlas also contains tissue from carotid artery, coronary artery, and femoral artery (Table S5, S6).

We trained CellTypist [15](v1.7.1) and scANVI [56] (scvi-tools v1.4.0.post1) on the same Tabula Sapiens 2.0 training cells as pretrained scBoost. We applied this pretrained CellTypist and scANVI model to every query dataset. We ran 10x Genomics cellranger annotate (v10.1.0) and Azimuth through the 10X Genomics Cloud Platform.

To compare scBoost with tested methods’ outputs, we compared accuracies using Wilcoxon signed-rank test at the donor level and we performed Holm-Bonferroni correction. The scBoost to tested method comparisons are not independent, and Holm-Bonferroni correction does not assume independence.

To transfer cell type labels from Wirka et al. [53] to the rest of the Plaque Atlas [50], we trained scBoost on Wirka et al. data in the Plaque Atlas (11,751 cells from coronary artery tissue, 4 donors, 10x 3’ v2), then applied it to the other 247,365 cells (69 donors, 11 studies, Table S6) without retraining. We trained two models: one on the level 1 labels (12 labels) and one on the level 2 labels (19 labels with at least 20 cells). Both followed the setup from the Tabula Sapiens 2.0 pretrained model except for two changes: 1) PCA and NMF used all genes detected in at least 3 training cells (18,848 genes), rather than 3,000 highly variable genes, and 2) there were four cross-validation folds that held one donor each. We reported the calls of the XGBoost meta-learner, focusing on the fibromyocyte, smooth muscle cell and fibroblast labels.

**Table S5:** Composition of the Plaque Atlas (Traeuble et al., Nat Commun 2025)

| Category | Number of Cells |
|---|---|
| Total | 259,116 |
| Assay: 10x 3′ v3 | 194,813 |
| Assay: 10x 3′ v2 | 64,303 |
| Sorting: Unsorted | 132,145 |
| Sorting: CD45<sup>+</sup> | 104,948 |
| Sorting: T cells only | 22,023 |
| Tissue: Carotid artery segment | 188,372 |
| Tissue: Coronary artery | 35,395 |
| Tissue: Femoral artery | 35,349 |

### scBoost comparison with popV

scBoost and popV are the only two methods, to our knowledge, that uses multiple established cell embedding or cell type classification methods as input to perform a final cell type prediction. The default for scBoost and popV are different, and this may create bias due to the performance of the individual input methods. To this end, we tested scBoost and popV using the same set of input cell embedding methods. popV’s default contains eight methods, four of which are cell embedding methods, which are applicable as input to scBoost. We selected all the cell embedding methods in the popV default for this test: scVI, scANVI, Harmony, and Scanorama.

For the Plaque Atlas and the fetal heart, we applied three pretrained popV [18] models from the Hugging Face Hub (All_Cells, Heart, and Vasculature), which were trained on Tabula Sapiens 2.0, the same reference as pretrained scBoost. popV All_Cells was applied to both datasets, popV Vasculature was applied to the Plaque Atlas, and popV Heart was applied to the Fetal Heart Map. We ran popV (v0.6.1) in its fast prediction mode, which applies the trained models without retraining six out of its eight member methods (CellTypist, kNN on scVI, OnClass, scANVI, a support vector machine and XGBoost). We ran popV with donor as the batch key, and we used its consensus prediction.

**Fig. S1:**
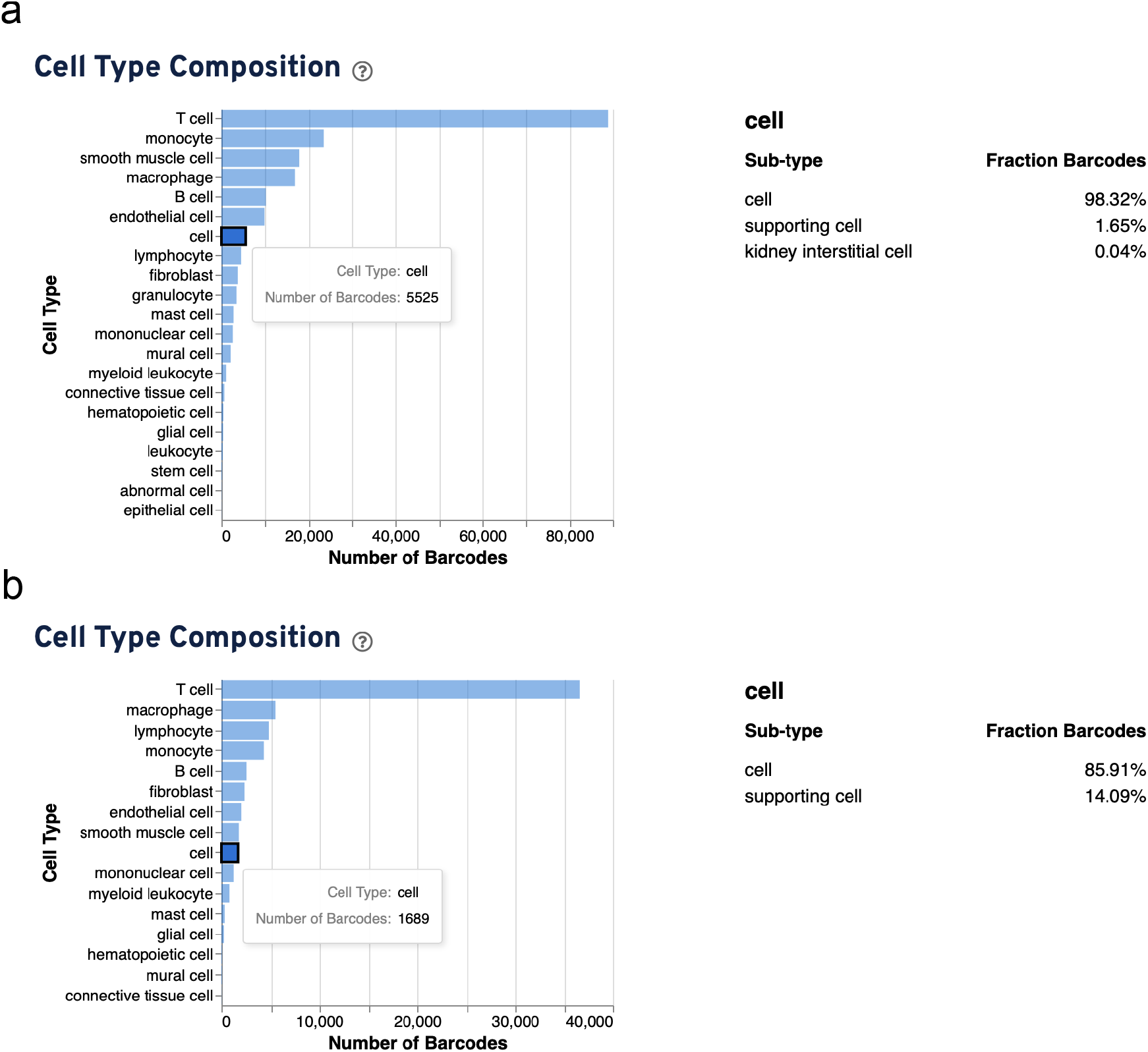
10X Genomics cellranger human plaque atlas annotation. **a**. 3’ v3 chemstry has 5,525 “cell” ambiguous annotation. **b**. 3’ v2 chemstry has 1,689 “cell” ambiguous annotation.

**Table S6:**
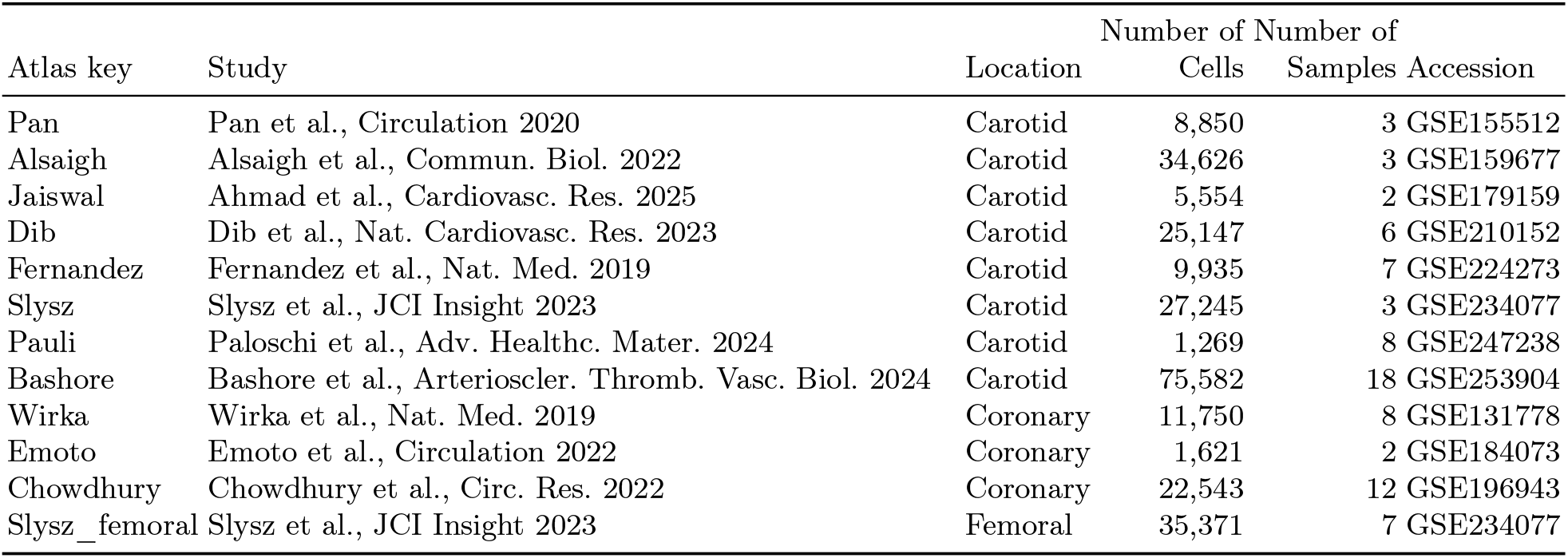
Source studies of the Plaque Atlas, from Table 1 of Traeuble et al., Nat Commun 2025.

|  |  |  | Number of | Number of |  |
|---|---|---|---|---|---|
| Atlas key | Study | Location | Cells | Samples | Accession |
| Pan | Pan et al., Circulation 2020 | Carotid | 8,850 | 3 | GSE155512 |
| Alsaigh | Alsaigh et al., Commun. Biol. 2022 | Carotid | 34,626 | 3 | GSE159677 |
| Jaiswal | Ahmad et al., Cardiovasc. Res. 2025 | Carotid | 5,554 | 2 | GSE179159 |
| Dib | Dib et al., Nat. Cardiovasc. Res. 2023 | Carotid | 25,147 | 6 | GSE210152 |
| Fernandez | Fernandez et al., Nat. Med. 2019 | Carotid | 9,935 | 7 | GSE224273 |
| Slysz | Slysz et al., JCI Insight 2023 | Carotid | 27,245 | 3 | GSE234077 |
| Pauli | Paloschi et al., Adv. Healthc. Mater. 2024 | Carotid | 1,269 | 8 | GSE247238 |
| Bashore | Bashore et al., Arterioscler. Thromb. Vasc. Biol. 2024 | Carotid | 75,582 | 18 | GSE253904 |
| Wirka | Wirka et al., Nat. Med. 2019 | Coronary | 11,750 | 8 | GSE131778 |
| Emoto | Emoto et al., Circulation 2022 | Coronary | 1,621 | 2 | GSE184073 |
| Chowdhury | Chowdhury et al., Circ. Res. 2022 | Coronary | 22,543 | 12 | GSE196943 |
| Slysz_femoral | Slysz et al., JCI Insight 2023 | Femoral | 35,371 | 7 | GSE234077 |

### Memory and Runtime Evaluation Methods

We used the Python memory-profiler package and wrapped it around the scBoost (ensemble) call to measure memory usage for four datasets (Tabula Sapiens, Human Immune Cell Atlas, GTEx, HypoMap) and across multiple input embedding dimensions (10, 25, 50, and 100 dimensions for each embedding method). We combined PCA, SCA, Harmony, Scanorama and scVI embeddings. We performed the evaluation on a machine with eight NVIDIA RTX A6000 GPUs and two AMD EPYC 7502 CPUs. We tracked run time via the Python time package and logged data loading, train/test split, fit, and prediction & evaluation stages time usage.

### scBoost-ensemble application to cell-type classification

To demonstrate scBoost’s application on cell-type classification task, we tested scBoost (ensemble) on Tabula Sapiens, GTEx[17], Hypomap[45], or Human Immune Cell Atlas[15] data with classic batch correction methods’ embeddings. We first generated embeddings using seven commonly-used embedding methods: PCA, SCA, Harmony, Scanorama, Geosketch with Scanorama, and scVI. We take these embeddings as inputs to scBoost-ensemble to train base classifiers for individual embedding method with out-of-fold (OOF) holdout, and we train meta-learners to combine the outputs across seven embedding methods’ base-learners. The best scBoost-ensemble model has 67.47% to 99.45% top-1 accuracy across the four datasets (Figure 1b, Table S12).

**Table S7:**
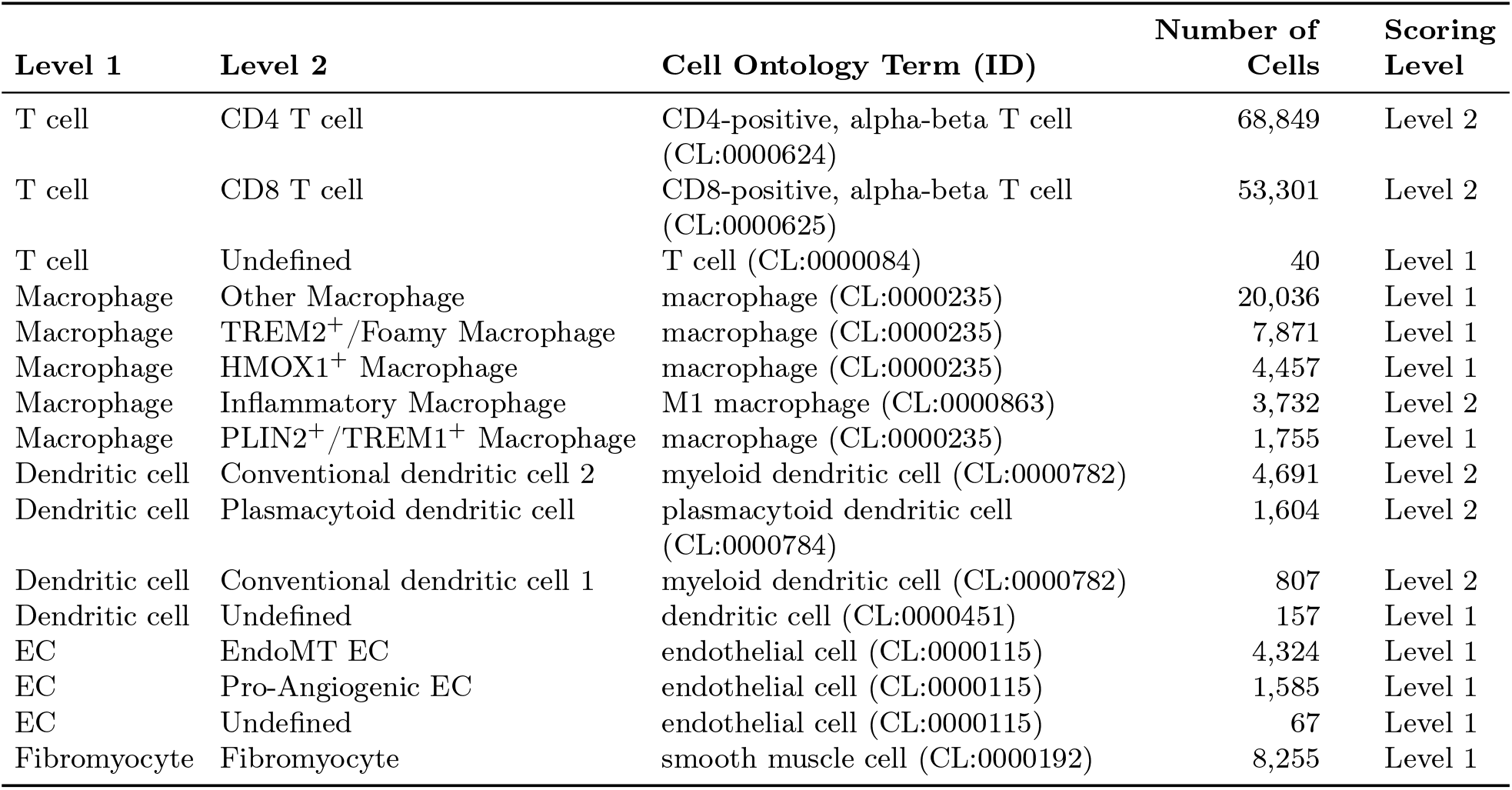
Manually matched Plaque Atlas authors’ level 1 and level 2 labels to the Cell Ontology term. If a level 2 label does not have a matching Cell Ontology term, we score it at the corresponding level 1 term.

| **Level 1** | **Level 2** | **Cell Ontology Term (ID)** | **Number of Cells** | **Scoring Level** |
|---|---|---|---|---|
| T cell | CD4 T cell | CD4-positive, alpha-beta T cell (CL:0000624) | 68,849 | Level 2 |
| T cell | CD8 T cell | CD8-positive, alpha-beta T cell (CL:0000625) | 53,301 | Level 2 |
| T cell | Undefined | T cell (CL:0000084) | 40 | Level 1 |
| Macrophage | Other Macrophage | macrophage (CL:0000235) | 20,036 | Level 1 |
| Macrophage | TREM2<sup>+</sup>/Foamy Macrophage | macrophage (CL:0000235) | 7,871 | Level 1 |
| Macrophage | HMOX1<sup>+</sup> Macrophage | macrophage (CL:0000235) | 4,457 | Level 1 |
| Macrophage | Inflammatory Macrophage | M1 macrophage (CL:0000863) | 3,732 | Level 2 |
| Macrophage | PLIN2<sup>+</sup>/TREM1<sup>+</sup> Macrophage | macrophage (CL:0000235) | 1,755 | Level 1 |
| Dendritic cell | Conventional dendritic cell 2 | myeloid dendritic cell (CL:0000782) | 4,691 | Level 2 |
| Dendritic cell | Plasmacytoid dendritic cell | plasmacytoid dendritic cell (CL:0000784) | 1,604 | Level 2 |
| Dendritic cell | Conventional dendritic cell 1 | myeloid dendritic cell (CL:0000782) | 807 | Level 2 |
| Dendritic cell | Undefined | dendritic cell (CL:0000451) | 157 | Level 1 |
| EC | EndoMT EC | endothelial cell (CL:0000115) | 4,324 | Level 1 |
| EC | Pro-Angiogenic EC | endothelial cell (CL:0000115) | 1,585 | Level 1 |
| EC | Undefined | endothelial cell (CL:0000115) | 67 | Level 1 |
| Fibromyocyte | Fibromyocyte | smooth muscle cell (CL:0000192) | 8,255 | Level 1 |

**Table S8:** Accuracy of the pretrained models the full Plaque Atlas (259,114 cells, 73 donors) and scored against the authors’ level 1 labels. Exact: cells prediction matches exactly to the Plaque Atlas’ authors’ labels. Same or finer (achievable): The model’s prediction matches to the same or finer term per Cell Ontology class, when the term also appeared in the pretrained set from Tabula Sapiens 2.0. *β* sets the class prior of the scBoost meta-learner (0: training class frequencies, 1: flat prior). Primary: the model setting that we use for comparison across methods. Holm p: exact Wilcoxon signed-rank test against pretrained scBoost (*β* = 0, xgboost meta) on per-donor Same or finer over the 73 donors, Holm-adjusted over 13 comparisons.

|  | **Trained on** |  |  |  |  |  |  |
|---|---|---|---|---|---|---|---|
| **Model** | **Dataset** | **Labels** | **Primary** | **Exact** | **Same or finer** | **Same, finer or coarser** | **Holm *p* vs scBoost** |
| scBoost pretrained, *β* = 0, xgboost meta | Tabula Sapiens 2.0 (637,267 cells) | 167 CL classes | yes | 0.439 | 0.887 | 0.887 | reference |
| scBoost pretrained, *β* = 0, logreg meta | Tabula Sapiens 2.0 (637,267 cells) | 167 CL classes |  | 0.434 | 0.892 | 0.892 | 1.2 × 10<sup>−5</sup> |
| scBoost pretrained, *β* = 0.5, xgboost meta | Tabula Sapiens 2.0 (637,267 cells) | 167 CL classes |  | 0.426 | 0.886 | 0.888 | 0.901 |
| scBoost pretrained, *β* = 0.5, logreg meta | Tabula Sapiens 2.0 (637,267 cells) | 167 CL classes |  | 0.414 | 0.887 | 0.893 | 0.840 |
| scBoost pretrained, *β* = 1, xgboost meta | Tabula Sapiens 2.0 (637,267 cells) | 167 CL classes |  | 0.372 | 0.841 | 0.850 | 3.6 × 10<sup>−17</sup> |
| scBoost pretrained, *β* = 1, logreg meta | Tabula Sapiens 2.0 (637,267 cells) | 167 CL classes |  | 0.314 | 0.794 | 0.876 | 2.8 × 10<sup>−21</sup> |
| scANVI, pretrained (scArches surgery) | Tabula Sapiens 2.0 (637,267 cells) | 167 CL classes | yes | 0.445 | 0.877 | 0.877 | 2.0 × 10<sup>−7</sup> |
| CellTypist, pretrained | Tabula Sapiens 2.0 (637,267 cells) | 167 CL classes | yes | 0.394 | 0.859 | 0.862 | 1.6 × 10<sup>−10</sup> |
| popV pretrained Vasculature, consensus | Tabula Sapiens 2.0, popV Vasculature (38,385 cells) | 25 classes | yes | 0.407 | 0.867 | 0.867 | 1.2 × 10<sup>−5</sup> |
| popV pretrained All_Cells, consensus | Tabula Sapiens 2.0, popV All_Cells (1,107,071 cells) | 179 classes | yes | 0.358 | 0.848 | 0.865 | 4.2 × 10<sup>−14</sup> |
| Azimuth pan-human, coarse | Pan-human Azimuth reference (23 tissues) | 85 classes (medium level) | yes | 0.725 | 0.787 | 0.796 | 7.2 × 10<sup>−19</sup> |
| Azimuth pan-human, fine | Pan-human Azimuth reference (23 tissues) | 291 classes (fine level) |  | 0.679 | 0.815 | 0.815 | 1.2 × 10<sup>−16</sup> |
| 10x Cloud, coarse | CELLxGENE Census | Cell Ontology terms (count not published) | yes | 0.777 | 0.755 | 0.852 | 3.0 × 10<sup>−20</sup> |
| 10x Cloud, fine | CELLxGENE Census | Cell Ontology terms (count not published) |  | 0.377 | 0.823 | 0.853 | 9.8 × 10<sup>−6</sup> |

**Table S9:** Accuracy of the pretrained models on the full Plaque Atlas (259,114 cells, 73 donors) against the authors’ level 2 labels, mapped to Cell Ontology terms by our reviewed label map. Exact: cells whose prediction matches the mapped term exactly. Same or finer (achievable): The model’s prediction matches to the same or finer term per Cell Ontology class, when the term also appeared in the pretrained set from Tabula Sapiens 2.0. *β* sets the class prior of the scBoost meta-learner (0: training class frequencies, 1: flat prior). Primary: the model setting that we use for comparison across methods. Holm p: exact Wilcoxon signed-rank test against pretrained scBoost (*β* = 0, xgboost meta) on per-donor Same or finer (achievable) accuracy over the 73 donors, Holm-adjusted over 13 comparisons.

|  | **Trained on** |  |  |  |  |  |  |  |
|---|---|---|---|---|---|---|---|---|
| **Model** | **Dataset** | **Labels** | **Primary** | **Exact** | **Same or finer (achievable)** | **Same or finer** | **Same, finer or coarser** | **Holm *p* vs scBoost** |
| scBoost pretrained, *β* = 0, xgboost meta | Tabula Sapiens 2.0 (637,267 cells) | 167 CL classes | yes | 0.792 | 0.794 | 0.783 | 0.802 | reference |
| scBoost pretrained, *β* = 0, logreg meta | Tabula Sapiens 2.0 (637,267 cells) | 167 CL classes |  | 0.800 | 0.804 | 0.793 | 0.807 | 1.2 × 10<sup>−8</sup> |
| scBoost pretrained, *β* = 0.5, xgboost meta | Tabula Sapiens 2.0 (637,267 cells) | 167 CL classes |  | 0.779 | 0.793 | 0.782 | 0.812 | 1.000 |
| scBoost pretrained, *β* = 0.5, logreg meta | Tabula Sapiens 2.0 (637,267 cells) | 167 CL classes |  | 0.776 | 0.796 | 0.785 | 0.808 | 1.000 |
| scBoost pretrained, *β* = 1, xgboost meta | Tabula Sapiens 2.0 (637,267 cells) | 167 CL classes |  | 0.653 | 0.713 | 0.703 | 0.763 | 6.7 × 10<sup>−17</sup> |
| scBoost pretrained, *β* = 1, logreg meta | Tabula Sapiens 2.0 (637,267 cells) | 167 CL classes |  | 0.613 | 0.678 | 0.668 | 0.789 | 1.1 × 10<sup>−20</sup> |
| scANVI, pretrained (scArches surgery) | Tabula Sapiens 2.0 (637,267 cells) | 167 CL classes | yes | 0.777 | 0.782 | 0.771 | 0.808 | 0.002 |
| CellTypist, pretrained | Tabula Sapiens 2.0 (637,267 cells) | 167 CL classes | yes | 0.746 | 0.757 | 0.746 | 0.755 | 3.3 × 10<sup>−8</sup> |
| popV pretrained Vasculature, consensus | Tabula Sapiens 2.0, popV Vasculature (38,385 cells) | 25 classes | yes | 0.763 | 0.770 | 0.758 | 0.769 | 1.6 × 10<sup>−5</sup> |
| popV pretrained All_Cells, consensus | Tabula Sapiens 2.0, popV All_Cells (1,107,071 cells) | 179 classes | yes | 0.669 | 0.733 | 0.723 | 0.761 | 3.7 × 10<sup>−17</sup> |
| Azimuth pan-human, coarse | Pan-human Azimuth reference (23 tissues) | 85 classes (medium level) | yes | 0.286 | 0.348 | 0.343 | 0.796 | 2.8 × 10<sup>−21</sup> |
| Azimuth pan-human, fine | Pan-human Azimuth reference (23 tissues) | 291 classes (fine level) |  | 0.252 | 0.381 | 0.378 | 0.812 | 2.8 × 10<sup>−21</sup> |
| 10x Cloud, coarse | CELLxGENE Census | Cell Ontology terms (count not published) | yes | 0.296 | 0.296 | 0.292 | 0.857 | 2.8 × 10<sup>−21</sup> |
| 10x Cloud, fine | CELLxGENE Census | Cell Ontology terms (count not published) |  | 0.390 | 0.608 | 0.599 | 0.748 | 3.7 × 10<sup>−17</sup> |

**Table S10:** Accuracy of the pretrained models on the Fetal Heart Map (633,954 nucli, 41 donors) where the authors’ label maps to a Cell Ontology term. Exact and Same or finer (achievable) are on the 520,909 nuclei whose term is one of the Tabula Sapiens 2.0 model’s classes. *β* sets the class prior of the scBoost meta-learner (0: training class frequencies, 1: flat prior). Primary: the model setting that we use for comparison across methods. Holm p: exact Wilcoxon signed-rank test against pretrained scBoost (*β* = 0, xgboost meta) on per-donor Same or finer (achievable) over the 41 donors, Holm-adjusted over 5 comparisons.

|  | **Trained on** |  |  |  |  |  |  |  |
|---|---|---|---|---|---|---|---|---|
| **Model** | **Dataset** | **Labels** | **Primary** | **Exact** | **Same or finer (achievable)** | **Same or finer** | **Same, finer or coarser** | **Holm *p* vs scBoost** |
| scBoost pretrained, β = 0, xgboost meta | Tabula Sapiens 2.0 (637,267 cells) | 167 CL classes | yes | 0.730 | 0.754 | 0.619 | 0.822 | reference |
| scBoost pretrained, β = 1, xgboost meta | Tabula Sapiens 2.0 (637,267 cells) | 167 CL classes |  | 0.701 | 0.727 | 0.597 | 0.751 | 4.9 × 10<sup>−6</sup> |
| scANVI, pretrained (scArches surgery) | Tabula Sapiens 2.0 (637,267 cells) | 167 CL classes | yes | 0.141 | 0.161 | 0.132 | 0.319 | 5.1 × 10<sup>−11</sup> |
| CellTypist, pretrained | Tabula Sapiens 2.0 (637,267 cells) | 167 CL classes | yes | 0.273 | 0.295 | 0.242 | 0.426 | 4.5 × 10<sup>−11</sup> |
| popV pretrained Heart, consensus | Tabula Sapiens 2.0, popV Heart (23,248 cells) | 22 classes | yes | 0.372 | 0.380 | 0.312 | 0.434 | 4.0 × 10<sup>−8</sup> |
| popV pretrained All_Cells, consensus | Tabula Sapiens 2.0, popV All_Cells (1,107,071 cells) | 179 classes | yes | 0.234 | 0.262 | 0.215 | 0.326 | 6.5 × 10<sup>−9</sup> |

**Table S11:** Memory usage (MB) for each dataset across embedding dimensionalities *N*_dim_.

|  | Memory usage (MB) |  |  |  |
|---|---|---|---|---|
| Dataset | *N*<sub>dim</sub> = 10 | *N*<sub>dim</sub> = 25 | *N*<sub>dim</sub> = 50 | *N*<sub>dim</sub> = 100 |
| Tabula Sapiens | 16 645.2 | 17 041.3 | 17 409.3 | 16 823.1 |
| Human Immune | 3950.6 | 3982.6 | 4031.6 | 4154.0 |
| GTEx | 7344.1 | 7373.4 | 7680.9 | 7690.3 |
| HypoMap | 10 442.2 | 10 496.7 | 10 626.0 | 10 889.2 |
| ICA | 11 739.5 | 11 962.7 | 12 081.7 | 12 241.0 |

**Table S12:** scBoost-ensemble cell-type classification results on four datasets. Each row is that architecture’s best configuration for that dataset over the same sweep (127 embedding subsets × 2 base learner settings (MLP, XGBoost) × 2 meta-learners (XGBoost, Elastic Net)); each architecture is therefore reported at its own optimum over an identical search space, rather than against a fixed-configuration baseline. Bold marks the best value in each dataset block. We display results from the donor hold-out split. The ensemble takes top-1 on the Immune Cell Atlas and ties on HypoMap, and takes top-5 on GTEx and Tabula Sapiens, where concatenation trails it by 6.4% of top-5 accuracy.

| Dataset | Architecture | Embeddings (*k*) | Top-1 | Top-5 | Macro-F1 |
|---|---|---|---|---|---|
| GTEx | (i) Best single embedding | Geosketch(+Scanorama) (1) | 0.9084 | 0.9832 | 0.7520 |
|  | (ii) scBoost-concat | Geosketch(+Scanorama),<br>PCA, Scanorama (3) | **0.9143** | 0.9770 | **0.7791** |
|  | (iii) scBoost-ensemble | Geosketch(+Scanorama),<br>Harmony, NMF,<br>Scanorama (4) | 0.9117 | **0.9900** | 0.7544 |
| Immune Cell Atlas | (i) Best single embedding | scVI (1) | 0.9073 | 0.9959 | 0.8751 |
|  | (ii) scBoost-concat | Geosketch(+Scanorama),<br>SCA, scVI (3) | 0.9144 | **0.9975** | 0.8872 |
|  | (iii) scBoost-ensemble | Harmony, PCA, Scanorama,<br>scVI (4) | **0.9167** | 0.9970 | **0.8882** |
| HypoMap | (i) Best single embedding | NMF (1) | **0.9945** | **0.9992** | **0.8931** |
|  | (ii) scBoost-concat | Geosketch(+Scanorama),<br>PCA, Scanorama (3) | 0.9938 | 0.9977 | 0.8903 |
|  | (iii) scBoost-ensemble | Geosketch(+Scanorama),<br>PCA (2) | **0.9945** | 0.9977 | 0.8930 |
| Tabula Sapiens | (i) Best single embedding | SCA (1) | 0.6530 | 0.9292 | 0.5056 |
|  | (ii) scBoost-concat | Geosketch(+Scanorama),<br>NMF, PCA, SCA,<br>Scanorama (5) | **0.6747** | 0.8748 | 0.5009 |
|  | (iii) scBoost-ensemble | SCA, scVI (2) | 0.6676 | **0.9391** | **0.5138** |

